# Reverse engineering Lewy bodies suggests α-synuclein pathology as condensatopathy

**DOI:** 10.64898/2026.10.09.757996

**Authors:** Roberto Sansevrino, Aleksandr Korobeinikov, Jian-Hua Chen, Christian Hoffmann, Hugo Le Roy, Kiersten M. Ruff, András Major, Asima Nayak, Paula Brosius, Axel Ekman, Shujin He, Michael Chen, Joshua Jackson, Gerard Aguilar Pérez, Han Wang, Johannes Vincent Tromm, Gwendolin Schneider, Kate Lau, Kathrin Textoris-Taube, Ludwig Sinn, Rinse de Boer, Lars T. Hofstede, Jonas Franz, Mark A Le Gros, Silvio O. Rizzoli, Tiago F. Outeiro, Paolo De Los Rios, Vadim Demichev, Michael Mülleder, Markus Ralser, Dietmar Schmitz, Daniele Bano, Geert van den Bogaart, Yifan Dai, Alberto J. Espay, Britta J. Eickholt, Christine Stadelmann, Pallavi P. Gopal, Rohit V. Pappu, Tim Salditt, Carolyn Larabell, Dragomir Milovanovic

## Abstract

Understanding how soluble monomeric α-synuclein transitions into pathological α-synuclein (Lewy bodies) has remained a central challenge in neurodegeneration. Existing models, largely based on treating cells and animals with pre-formed α-synuclein fibrils cannot account for the emergence of the first inclusion without pre-existing seeds. Here, we combine in vitro reconstitutions, computational modeling and a minimal cellular system to show that phase separation of α-synuclein with an interacting scaffold can recapitulate Lewy body-like structures. An unbiased proteomics screen, live-cell imaging, correlative light-electron microscopy, and soft X-ray tomography show that Lewy body-like structures are associated with redistribution of actin and membrane-bound organelles, with mitochondria accumulating at their periphery in an actin-dependent manner. Structural analyses further indicate that α-synuclein condensates provide local environments in which the insoluble (-sheets can form. Together, these findings support a seed-independent trajectory from soluble α-synuclein through condensation and progressive immobilization to Lewy body-like assemblies and, subsequently, insoluble (-sheet fibrils. This framework offers a new mechanistic route for understanding the disease progression and promises to guide the design of innovative therapeutic interventions.

## INTRODUCTION

The presence of proteinaceous inclusions, referred to as Lewy bodies (LBs), is a defining histological signature of Parkinson’s disease (PD) and related disorders^1,2^. LBs are abnormal protein deposits that accumulate inside neurons. These deposits primarily contain α-synuclein^3^, are a pathological feature of synucleinopathies^2^, and can also be detected in tissues from patients with Alzheimer’s disease related dementias^4^. Classically, LBs were considered to be nearly exclusively formed by amyloid-like structures made of α-synuclein and other misfolded proteins.

Currently available animal and cellular models all require preformed α-synuclein fibrils to trigger the α-synuclein lesions^5^. However, several recent lines of evidence challenge the view of lesions being scaffolded by fibrillar structures. For example, only a small fraction of LBs contains α-synuclein fibrils^6,7^. Moreover, while α-synuclein fibrils have a well-defined structure^8,9^, α-synuclein deposits within the LBs exhibit different morphologies^10^. Even though α-synuclein is a key component of LBs, immunohistochemical studies^11,12^ and mass spectrometry suggest that LBs isolated from PD patients include nearly three hundred different proteins^13,14^ such as mitochondrial proteins, components of the ubiquitin–proteasome system and proteins involved in autophagy and aggresome formation. Furthermore, many of these proteins (e.g., DJ-1, LRRK2, parkin, PINK-1) have been implicated in the pathogenesis of inherited PD^15^. Postmortem analysis of patients with dementia with LBs further indicated that these inclusions are in fact a complex medley packed with fragmented membranes, vesicular structures, and dysmorphic organelles (mitochondria, lysosomes, and autophagosomes)^16^. In particular, mitochondria tend to accumulate at the periphery of the LB inclusions^16,17^. Ultrastructural cryo-EM analyses of post-mortem LB disorders have shown a common Lewy filament core across the examined PD, Parkinson’s disease dementia, and dementia with Lewy bodies samples^18^, indicating that LBs provide a distinct microenvironment that may help catalyze the formation of fibrillar structures.

Protein condensation and aggregation may not necessarily constitute the initiating disease process. They may represent reactive transitions induced by preceding toxic, metabolic, inflammatory, infectious, genetic, or other biological stressors that alter protein concentration, solubility, post-translational modification, or cellular clearance. Under this interpretation, LB formation could mark a cellular response to an upstream disturbance rather than serve as its vehicle of propagation or primary cause of neuronal injury.

Here, we examine whether LB-like bodies can emerge through phase separation of α-synuclein with interacting scaffold proteins, accompanied by redistribution of actin and intracellular membranes of secretory vesicles and mitochondria^19–21^. Using a minimal reconstitution system of synphilin-1 as the scaffold protein for phase separation we recapitulated the key molecular features of LBs including them being a reaction center that appears to catalyze the nucleation of α-synuclein cross-β-sheet fibrils. Our successful reconstitution of LBs using principles of condensate biology enables us to investigate a seed-independent route to inclusion formation without presupposing whether the resulting structures are toxic, protective, or markers of the depletion of physiologically available soluble proteins. It allows us to rethink strategies for the design of therapeutics by opening up avenues that go beyond targeting individual molecules to modulating the viscoelastic and electrochemical properties of condensates.

## RESULTS

### Synphilin-1 and α-synuclein recapitulate Lewy body-like morphology

We systematically reviewed proteins reported to accumulate in LBs and focused on synphilin- 1 – a presynaptic protein that binds to synaptic vesicles^22^ and contains multiple scaffolding and coiled-coil domains that allow its interaction with many proteins, including α-synuclein^23–25^. In addition, synphilin-1 contains a large intrinsically disordered region (IDR). Together, these features suggest that synphilin-1 might be driver of phase separation. To test this, we purified recombinant EGFP-synphilin-1 (**Supplementary Fig. 1A, 1C**) and assessed its ability to undergo phase separation *in vitro* (**Fig. 1A**). We reconstituted different concentrations (1 µM, 5 µM, 10 µM) of protein with increasing percentages of polyethylene glycol 8000 (PEG 8,000; 0%, 0.5%, 1%, 3%, 5%, 10%) in 25 mM Tris-HCl (pH 7.4), 300 mM NaCl, 0.5 mM TCEP and incubated for 5 min at room temperature (RT). EGFP-synphilin-1 underwent phase separation at low concentrations and low percentages of crowding agent, as also schematized in the phase diagram (**Fig. 1B; Supplementary Fig. 1E**).

**Fig. 1.**
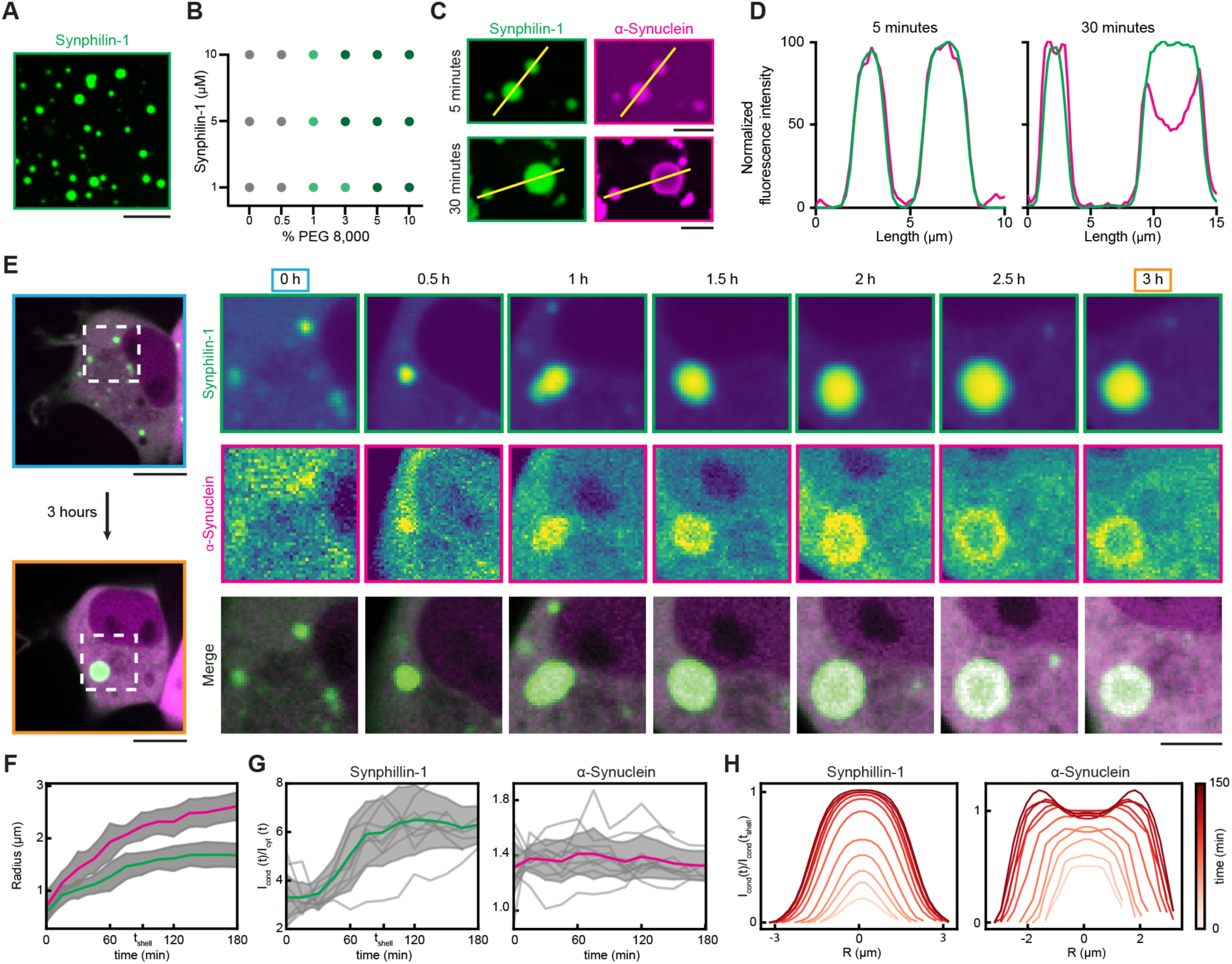
α-Synuclein demixes from synphilin-1 condensates forming a core-shell structure reminiscent of the Lewy body architecture. **(A)** Purified EGFP-synphilin-1 (5 µM) forms condensates in 3% PEG 8,000 at room temperature. Scale bar, 10 µm. **(B)** Mapping of the combinations of EGFP-synphilin-1 and PEG 8000 where phase separation was observed (green circles) and not observed (grey circles). Phase separation was determined as the appearance of droplets at the coverslip surface 5 min after the beginning of the incubation. Gray circles: no condensates; light green indicate the formation of condensates smaller than 1 µm^2^; dark green: condensates larger than 1 µm^2^. Each condition was assessed by three independent reconstitutions. **(C)** Reconstituted condensates containing EGFP-synphilin-1 (10 µM) and α-synuclein (20 µM) in 10% PEG 8,000 at ambient temperature 5 min (top) and 30 min (bottom) upon incubation. Note that both proteins colocalize in condensates after 5 min and demix to reach the core/shell structure after 30 min. Scale bar, 5 µm. **(D)** Line profiles of the highlighted selections in *C.* **(E)** Representative images of cells expressing EGFP-synphilin-1 and α-synuclein-BFP. Cell overview (left) and magnified representative images from indicated time points (right). Live-cell imaging indicates the growth of condensates over the period of 3 h, starting from the first mesoscale (>0,5 µm) puncta appearance. Note the demixing of α-synuclein and the evolution of the core-shell morphology. Scale bars, 10 µm (overview), 5 µm (magnified). **(F)** Quantification of the condensate radius over time for synphilin-1 (green line) and α-synuclein (magenta line), highlighting the larger size increase of the α-synuclein shell comparing to the synphilin-1 core. t_shell_ marks the time at which the third phase appear, concomitant with the plateau of relative intensity. **(G)** Variation in the relative concentration within condensates compared to the cytoplasm for synphilin-1 (left) and α-synuclein (right). I_cond_ and I_cyt_ stand for the intensities within the condensate and the cytoplasm, respectively. t_shell_ marks the time at which the third phase appear, concomitant with the plateau of relative intensity. Each gray trajectory denotes a distinct cell, with the colored line showcasing the mean and the shaded area indicating a range of two standard deviations. Data of ten condensates, each from a different cell, from three independent replicates. **(H)** Radial fluorescence intensity distributions (i.e., cross-sections averaged in all directions) from the central point of the condensate for synphilin-1 (left) and α-synuclein (right). The spatial coordinates of the condensate are normalized by its radius. The color code of the scheme reflects the time when each cut was taken. Note that as time progresses, the size of the condensate increases.

Since α-synuclein is well-established interaction partner of synphilin-1^23^, we sought to assess their putative co-assembly into condensates. *In vitro* reconstitutions using purified synphilin-1 and chemically labeled α-synuclein demonstrated a peculiar time-dependent demixing behavior. Specifically, at an initial stage (5 min incubation), α-synuclein was homogeneously enriched in synphilin-1 condensates, whereas over a 30 min time period a remodeling was observed with the condensate forming a core-shell morphology comprising a synphilin-1 core and an the α-synuclein shell. (**Fig. 1C, 1D**). Of note, α-synuclein on its own could undergo phase separation only in the presence of a high percentage of PEG 8,000 (10% PEG or higher) or when present in high concentrations (75 μM)^26–28^ (**Supplementary Fig. 1F, 1G**). These are likely to be non-physiological levels of crowders and protein concentrations.

Next, to test whether the minimal system of two proteins remains functional in a complex cellular environment, we ectopically expressed the proteins in non-neuronal cells. Overexpression of EGFP-synphilin-1 alone in HEK293 cells was sufficient to potentiate the formation of highly mobile inclusions of disparate sizes (**Supplementary Fig. 2A**). Following 48 h post-transfection, the condensates showed signs of undergoing physical aging transitions as evidenced by an increase in the immobile fraction in fluorescence recovery after photobleaching (FRAP) experiments (**Supplementary Fig. 2B, 2C**). Synphilin-1-containing condensates could undergo multiple events of fusion that led to larger condensates over time (**Supplementary Fig. 2D**). However, the circularity of condensates dropped over time, suggesting that synphilin-1 condensates either become dynamically arrested or convert to solid phases^29–31^ (**Supplementary Fig. 2E**).

Co-expression of EGFP-synphilin-1 and α-synuclein-BFP in HEK293 cells led to the formation of micron-scale condensates with core-shell morphology, recapitulating *in vitro* findings (**Fig. 1E**). We further analyzed the mobility of molecules inside these multi-phase condensates using FRAP and focusing on the mobility of synphilin-1 and α-synuclein. While EGFP-synphilin-1 alone was highly mobile at the rim of condensates and showed fluorescence recovery of about 50% (**Supplementary Fig. 2B, 2C**), its mobility dramatically decreased when co-transfected with α-synuclein-BFP, showing a fluorescence intensity recovery of only 20% (**Supplementary Fig. 2F, 2G**). These results indicate that α-synuclein decreases the mobility of synphilin-1, and that this reduced mobility comes from strong and complementary interactions between the core and shell layers.

The evolution from homogeneous condensates to a core-shell morphology could be under thermodynamic or nonequilibrium/dynamical control^30,32^. To distinguish between these two scenarios, we used fluorescence image intensity as an indicator to assess the relative concentrations of different cell regions and across different time points (**Fig. 1E, Supplementary Fig. 3A, 3B**). Initially, the linear increase in total cell protein concentration translated into the enrichment of proteins in the condensates (**Fig. 1F, Supplementary Fig. 3B**). While the α-synuclein concentration increases at a steady rate, its demixing begins only after the threshold concentration of synphilin-1 is reached within condensates (**Fig. 1G**). Once the shell starts emerging, the concentration of proteins in the condensates stays fixed. Interestingly, α-synuclein forms a fluid shell around the synphilin core (**Fig. 1H, Supplementary Movie 1**), reminiscent of condensate demixing observed in the nucleus^33,34^. Consistent with the equilibrium theory of phase separation in ternary mixtures^35^, the emergence of a third α-synuclein-rich phase that coexists with the synphilin-1-rich and dilute phases, fixes the concentration of protein in each phase. Thus, additional protein synthesis solely translates into a condensate volume increase (**Fig. 1H, Supplementary Fig. 3C**). Importantly, we propose that the mesoscale features of condensates bear resemblance to the morphology of Lewy bodies observed in Parkinson’s disease brain tissue^23^. Accordingly, we refer to the condensates being LB-Like inclusions (LBLs).

To assess the differences in material/structural properties between the core and the shell of LB-like inclusions, we turned to soft X-ray tomography (SXT). SXT is a cutting-edge isotropic 3D imaging method that requires no additional labeling or sophisticated chemical processing of the samples^36,37^. Moreover, the contrast in SXT images arises from the absorption by the bio-organic matter containing carbon and nitrogen from the intracellular organelles. The absorption differences between water and organelles yield a unique linear absorption coefficient (LAC)^37^. LACs allow for discrimination between similar organelles, such as lysosomes, mitochondria, or a cohort of viral particles in the cytosol^38^, as well as the densely packed structures such as chromatin territories^39^.

Indeed, in our tomograms we were able to identify both synphilin-1 condensates (**Fig. 2A-2D**) and LBLs **(Fig. 2E-2H)**. LBLs but not synphilin-1-only condensates exhibited the core-shell pattern based on their LAC differences (**Fig. 2A-2H**). In the SXT-reconstructed virtual slices, we readily observed the fusion of multiple LBLs with one another (**Fig. 2I, 2J; Supplementary Fig. 4; Supplementary Movie 2**). In line with our FRAP measurements, SXT tomograms 24 and 48 h post-transfection show higher molecular density of both synphilin-1 condensates and LBLs at the later time point (**Supplementary Fig. 5**), which is indicative of condensate coarsening as a function of time^40^.

**Fig. 2.**
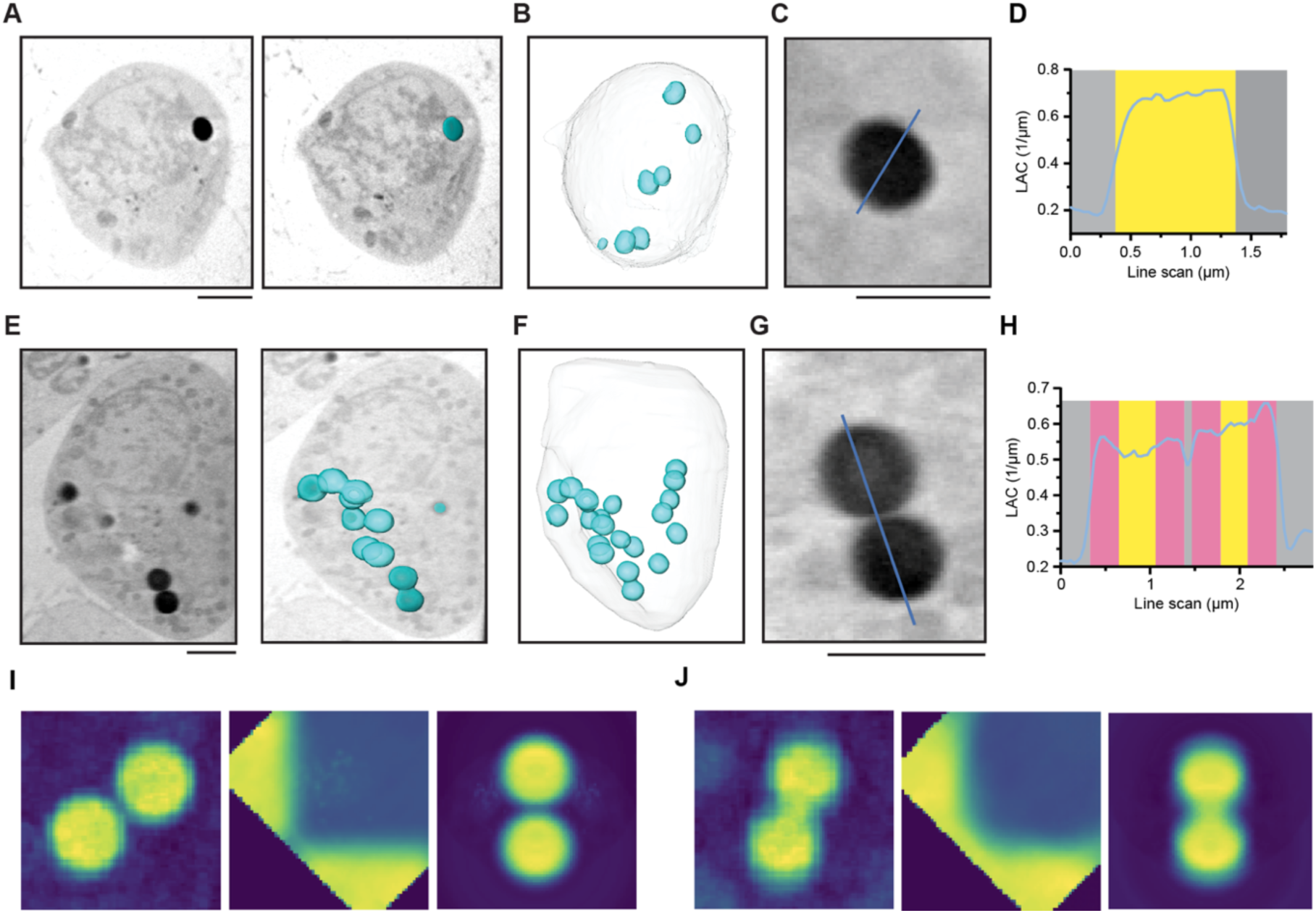
Soft X-ray tomography (SXT) of Lewy body-like condensates. **(A)** Orthoslice (virtual section) from a representative tomogram of a cell expressing EGFP-synphilin-1. Left, single slice of an X-ray tomogram; right, highlighted surface-rendered synphilin-1 condensate (cyan). Scale bar, 3 µm. **(B)** Surface-rendered view of the cell in *A*, highlighting all synphilin-1 condensates inside the cell. (**C)** Virtual section of a synphilin-1 condensate. Scale bar, 2 µm. **(D)** Line profile through the equatorial plane indicates a homogeneous linear absorption coefficient (LAC) of contents. **(E)** Virtual section from representative tomogram of a cell expressing LBL condensates (EGFP-synphilin-1 and α-synuclein-BFP). Left, single orthoslice of an X-ray tomogram; right, highlighted surface-rendered LBLs (cyan). Scale bar, 3 µm. **(F)** Surface-rendered cell in *E*, highlighting all LBLs inside the cell. **(G)** Virtual section of two LBLs. Scale bar, 2 µm. **(H)** Line profile through the equatorial plane showing the changes in LAC. Note the distinct absorption pattern at the core (yellow) and shell (magenta) of the condensate, indicating distinct material properties of these two regions within a condensate. **(I)** Left: orthoslice of a pair of LBLs in close proximity but remaining non-fused (i.e., the absence of content mixing); middle: 2D histogram of LAC values as a function of radial distance; right: representative interpolation of the LBL pair. **(J)** The same as in *I* but for LBLs that undergo fusion-fission (i.e., the presence of content mixing).

The time-dependent progression from apparently homogeneous condensates to core-shell architecture appears to be under thermodynamic control. This would suggest that the two structures represent distinct regions on the ternary phase diagram defined by activities of synphilin-1, α-synuclein, and the cellular milieu. To test for this possibility, we disrupted the ionic strength by exchanging the growth media of cells with solutions of different osmolarity. First, we replaced the medium with a hypotonic solution (DMEM diluted 20 times in H_2_O), knowing that local concentration and ionic strength are two important features that affect solubility of proteins and, by consequence, their ability to undergo phase separation. The application of hypotonic solution to cultured cells decreases the extracellular concentration of salts, resulting in a decrease in osmotic pressure inside the cell, which chases an uptake of water and the swelling of cells. This leads to a decrease of the bulk protein concentration^41^. In contrast, application of hypertonic solution to cultured cells resulted in an increase of extracellular salt concentration, leading the cell to lose water due to increased osmotic pressure and causing cell to shrink thereby increasing the bulk protein concentration. Small and highly dynamic condensates of EGFP-synphilin-1 fully dissolved after exchange with the hypotonic solution, while large condensates persisted (**Supplementary Fig. 6A**). Application of the hypertonic solution on cells expressing EGFP-synphilin-1 promoted the effective condensation as suggested by the appearance of a higher number of new fluorescence foci (**Supplementary Fig. 6B**). In cells containing LBLs, α-synuclein fully mixed with synphilin-1 condensates when stressed by hypotonic solution (**Supplementary Fig. 6C**). In contrast, LBLs kept maintained their core-shell architecture when stressed by hypertonic solution (**Supplementary Fig. 6D**). Together, these findings suggest that the distinct morphologies represent distinct stable states, albeit in different regions of the ternary phase diagram.

### Molecular grammars of α-synuclein encode Lewy body-like structures

Molecular grammars of IDRs, defined by non-random compositional biases and non-random patterning of different residues types with respect to one another, are linked with their functional properties and cellular localization^42–44^. We used NARDINI+ to uncover the molecular grammars of synphilin-1 and α-synuclein IDRs (**Fig. 3 A-3C**). This analysis revealed that α-synuclein has a high non-random hydrophobicity arising mainly from its hydrophobic region but also influenced by hydrophobic blocks in the amphipathic region. In α-synuclein, we also observed a non-random clustering of basic residues, and this overlaps with the amphipathic region. The grammar within synphilin-1 IDRs is complementary whereby hydrophobic and acidic residues are non-randomly distributed into well-defined linear clusters.

**Fig. 3.**
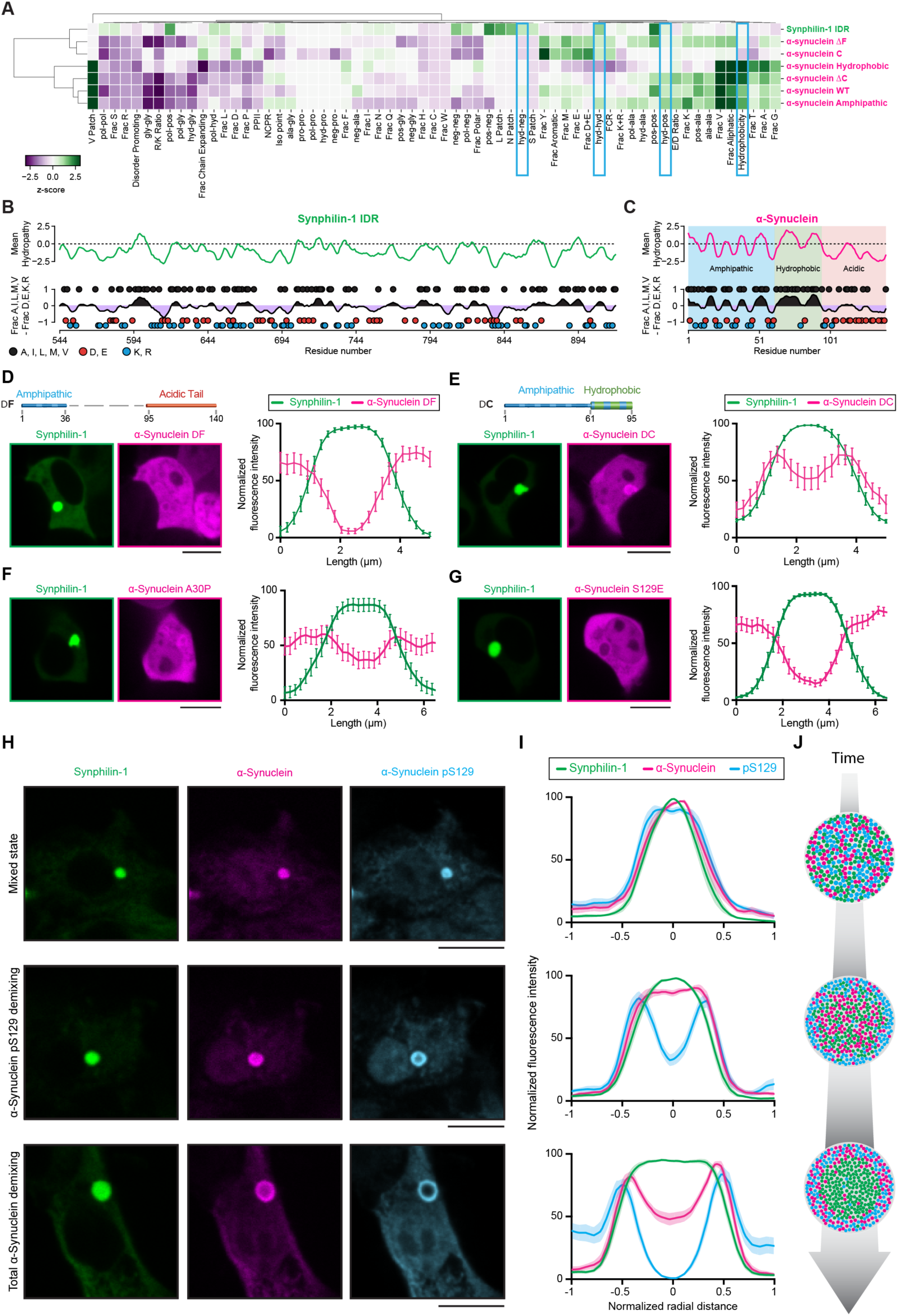
Molecular Grammar of Lewy body-like condensates. **(A)** Cluster map of NARDINI+ z-score vectors highlights distinctive molecular grammar features of synphilin-1 IDR and α-synuclein variants. **(B)** Schematic showing synphilin-1 IDR mean hydropathy score (top) and mean fraction of aliphatic residues minus fraction of charged residues (bottom). **(C)** Same schematic as in *B* for α-synuclein. **(D)** Representative images (left) of cells 24 h after co-transfection with a truncated version of α-synuclein-BFP (deleted residues Δ37-94, so-called ΔF; top scheme) and EGFP-synphilin-1 and cumulative line profile (right), indicating the demixing of α-synuclein ΔF from synphilin-1. Scale bar, 10 µm. **(E)** Representative images (left) of cells 24 h after co-transfection with a truncated version of α-synuclein-BFP (deleted residues Δ96-140, so-called ΔC; top scheme) and EGFP-synphilin-1 and cumulative line profile (right) indicating the demixing of α-synuclein ΔC from synphilin-1. Scale bar, 10 µm. **(F)** Representative images (left) of cells 24 h after co-transfection with α-synuclein-BFP A30P and EGFP-synphilin-1. and cumulative line profile (right) indicating the demixing of α-synuclein A30P from synphilin-1. Scale bar, 10 µm **(G)** Representative images (left) of cells 24 h after co-transfection with S129E phosphomimetic version of α-synuclein-BFP and EGFP-synphilin-1 and cumulative line profile (right) indicating the demixing of α-synuclein S129E from synphilin-1. Scale bar, 10 µm. **(H)** Representative images of HEK293 cells expressing early-stage LBLs (top; EGFP-synphilin-1 (green) and α-synuclein-BFP (magenta) are fully mixed), intermediate (middle), and late-stage LBLs (bottom; EGFP-synphilin-1 and α-synuclein-BFP form core-shell morphology) and immunostained for pS129 α-synuclein (blue). Note that pS129 α-synuclein starts accumulating at the periphery of condensates at an earlier stage when the total, unphosphorylated α-synuclein still remains enriched across the entire condensate. Scale bar, 10 μm. **(I)** The cumulative line profiles of conditions in *H* showing progressive pS129 and total α-synuclein demixing. **(J)** Schematic representation recapitulating the maturation of LBLs. At early stages, all components are intermixed; as the condensates mature, the internal rearrangements favor the demixing of α-synuclein in the shell and the enrichment of pS129 α- synuclein. The data in D-G and I represent an average ± s.e.m. of ten condensates each from a different cell from three independent replicates/experiments.

To test the impacts of distinctive grammars on the formation of the core-shell structure of LBLs, we prepared a series of mutants, including two truncated versions of α-synuclein: (i) α-synuclein(ΔF)-BFP, which lacks the central hydrophobic NAC domain (Δ 37–94 aa) responsible for the formation of fibrils^45^ and (ii) α-synuclein(ΔC)-BFP, which lacks the acidic C-terminal tail (Δ 96-140 aa). Furthermore, we assessed the effects of a point mutation, α-synuclein A30P, which is a common mutation identified in Parkinson’s disease patients^46^. We also assessed the impact of the S129 phosphomimetic substitution (S129E), as this site is traditionally found phosphorylated in patient Lewy bodies^47,48^.

While the co-expression of EGFP-synphilin-1 and α-synuclein(ΔF)-BFP in cells promoted the formation of canonical LBLs similar to those observed with the wild-type α-synuclein (**Fig. 3D**), co-expression of EGFP-synphilin-1 and α-synuclein(ΔC)-BFP caused only a partial de-mixing of the two proteins, with a higher frequency of condensates being homogeneously enriched in both proteins (**Fig. 3E**). α-synuclein A30P demonstrated significantly lower demixing in comparison with the wild-type (**Fig. 3F**).

For the formation of LBLs, the S129E mutant remained highly demixed, suggesting that a simple addition of negative charge to already highly acidic tail of α-synuclein has no major impact (**Fig. 3G**). However, to understand the actual phosphorylation pattern in LBLs, we immunostained transfected cells with antibodies against pS129 α-synuclein. These stainings showed progressive demixing of pS129 α-synuclein to the shell over time (**Fig. 3H-3J**), forming structures strikingly similar to LBs found in patients^10^. It is also worth noting that pS129 fraction demixed earlier than the total α-synuclein, suggesting that timing of protein accumulation and spatial organization of cellular components may play a particularly important role in triggering the formation of LBs.

### Lewy body-like condensates sequester actin

To examine how the cell responds to the appearance of LBs, we performed an unbiased proteomics screen of cells expressing LBLs. For this, we created AK-11 – an inducible α-synuclein-Halo stable cell line (**Supplementary Fig. 7**) – and transfected it with EGFP-synphilin-1 simultaneously with induction of α-synuclein expression. Cells were collected 24 h after transfection, subjected to downstream sample processing, and analyzed by LC MS-MS. In contrast to untransfected and uninduced cells, we observed broad differences in the cellular proteome involving cytoskeletal, mitochondrial, endolysosomal, plasma membrane, nuclear, endoplasmic reticulum, and Golgi-associated proteins **(Fig. 4A)**. Because these measurements compare inclusion-expressing cells with untransfected and uninduced controls, they do not distinguish direct effects of inclusions and responses to protein overexpression, redistribution of proteins into condensates, or depletion from their soluble functional pools.

**Fig. 4.**
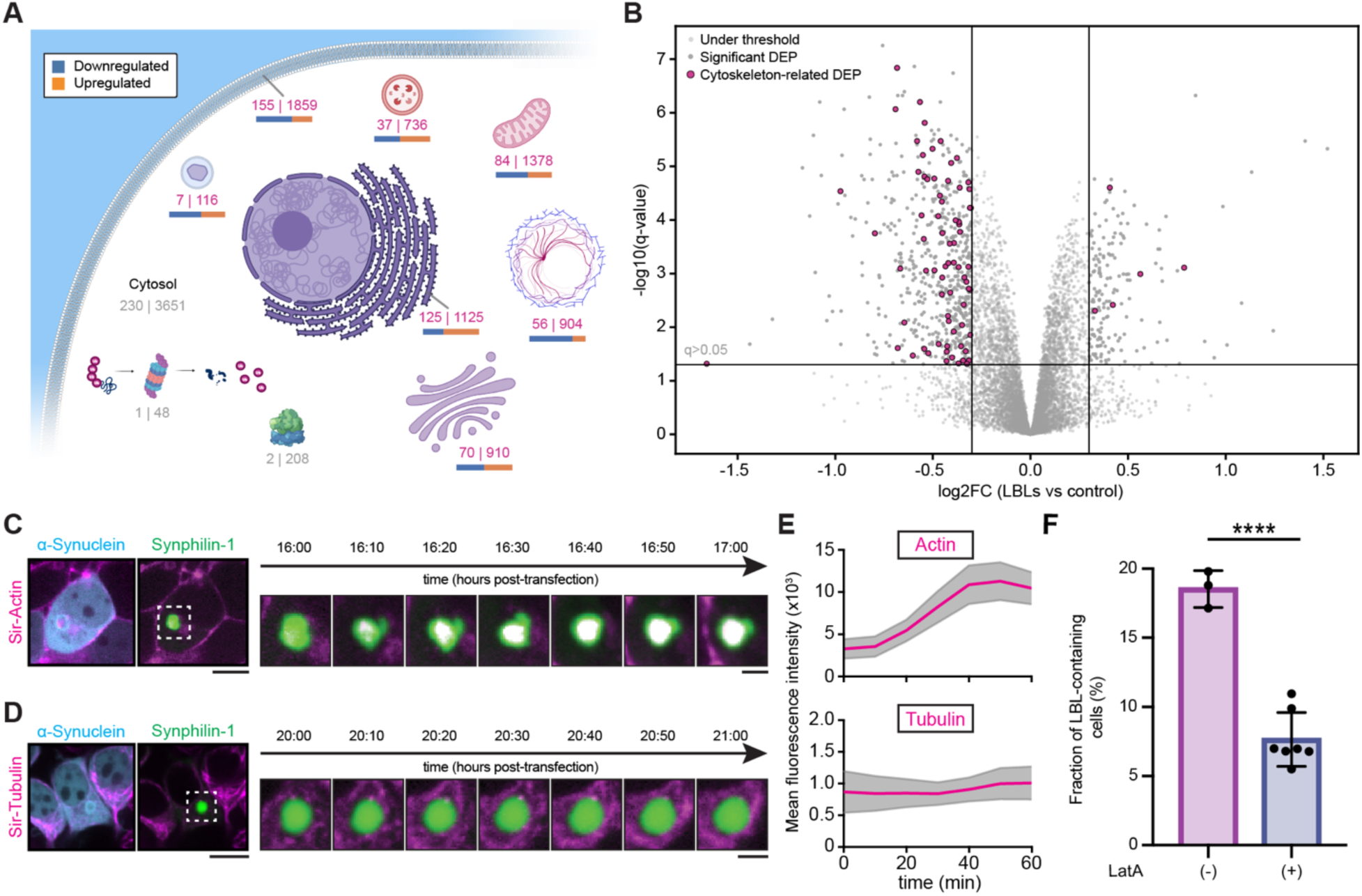
Lewy body-like condensates are associated with actin-dependent mitochondrial redistribution and altered morphology. **(A)** The proteome of LBLs-expressing cells is altered across a wide range of subcellular compartments. Numbers represent the number of significantly changed proteins (left) and total number of detected proteins (right) for each compartment. Protein subcellular localization was determined by Gene Ontology: Cellular Compartment analysis. Significance threshold: q≥0.05, |log2FC|≥0.3. **(B)** The volcano plot demonstrates changes in cytoskeletal proteins expression in LBLs-expressing cells. Magenta dots represent significantly changed cytoskeleton-related proteins. Significance threshold: q≥0.05, |log2FC|≥0.3. **(C)** Live-cell imaging demonstrates the enrichment of SiR-Actin inside the condensates (green, EGFP-synphilin-1; magenta, SiR-Actin; α-synuclein is not shown for clarity). Left, whole-cell overview; right, magnified condensate. Scale bar, 10 µm. **(D)** Live-cell imaging indicates no accumulation of SiR-Tubulin inside the condensates (green, EGFP-synphilin-1; magenta, SiR-Tubulin; α-synuclein is not shown for clarity). Left, whole-cell overview; right, magnified condensate. Scale bar, 3 µm. **(E)** Quantification of SiR-Actin (top) and SiR-Tubulin (bottom) signal inside the condensates (three independent experiments, fifteen condensates analyzed for each reporter). Note the enrichment of actin signal inside the condensate and the steady levels (i.e., the absence) of SiR-Tubulin signal. **(F)** Latrunculin A treatment inhibits formation of LBLs in AK-11 cells, expressing EGFP-synphilin-1 and α- synuclein-Halo in comparison with untreated controls, as represented by the fraction of LBLs-containing cells. Data from at least 2000 cells across at least three independent replicates for each condition. Error bars represent SD; dots represent individual replicates. Significance was tested using a Student’s t-test; asterisks, significance: ****p<0.0001.

Lewy bodies are known to contain cytoskeletal elements^10,16^, and expression of many cytoskeletal proteins was altered in cells that feature LBLs **(Fig. 4B)**, which prompted us to investigate how the cytoskeleton affects the formation of LBLs. To characterize this, we turned to live-cell imaging of condensate maturation over time. Using fluorescent actin or tubulin probes (SiR-Actin or SiR-Tubulin, respectively) and fluorescently labeled LBLs, we were able to simultaneously image the growth of LBLs and cytoskeleton dynamics. Imaging revealed that actin but not tubulin (**Fig. 4C-4E**) specifically enriched in LBLs over time.

To investigate whether LBLs formation is actin-dependent, we treated cells with 2.5 µM Latrunculin A (LatA, an actin polymerization inhibitor) after induction of LBLs expression. Cells treated with LatA demonstrated almost 3-fold decrease in LBLs formation frequency in comparison with a DMSO control, suggesting that filamenous actin is not only sequestered to LBLs, but is also required for their formation (**Fig. 4F**).

### Ultrastructural analyses of membrane-bound organelles surrounding α-synuclein inclusions

One of the morphological features of LBs is the accumulation of membrane-bound organelles in their near proximity. Whether this represents active sequestration by the inclusion, impaired trafficking, or a cellular containment response is unresolved. Given that the functional actin network is central to the intracellular dynamics and positioning of mitochondria^49–51^, combined with our observed actin-sequestering properties of LBLs, we investigated the behavior of membrane-bound organelles in LBLs-expressing cells. We employed FM4-64, LysoTracker-647 and MitoTracker-647 dyes to detect membrane infiltration, lysosomes and mitochondria, respectively. Both synphilin-1 condensates and LBLs appear to have a local enrichment of lipids at their interface (**Supplementary Fig. 8**). Interestingly, while the mitochondria in the proximity of synphilin-1 condensates formed characteristic elongated structures, they appeared to be more fragmented in the vicinity of LBLs (**Fig 5A, Supplementary Fig. 8A**). Correlative light-electron microscopy further confirmed the presence of membrane-bound organelles, such as mitochondria and ER, in close proximity and at the interface of condensates (**Fig. 5B, Supplementary Fig. 9**).

**Fig. 5.**
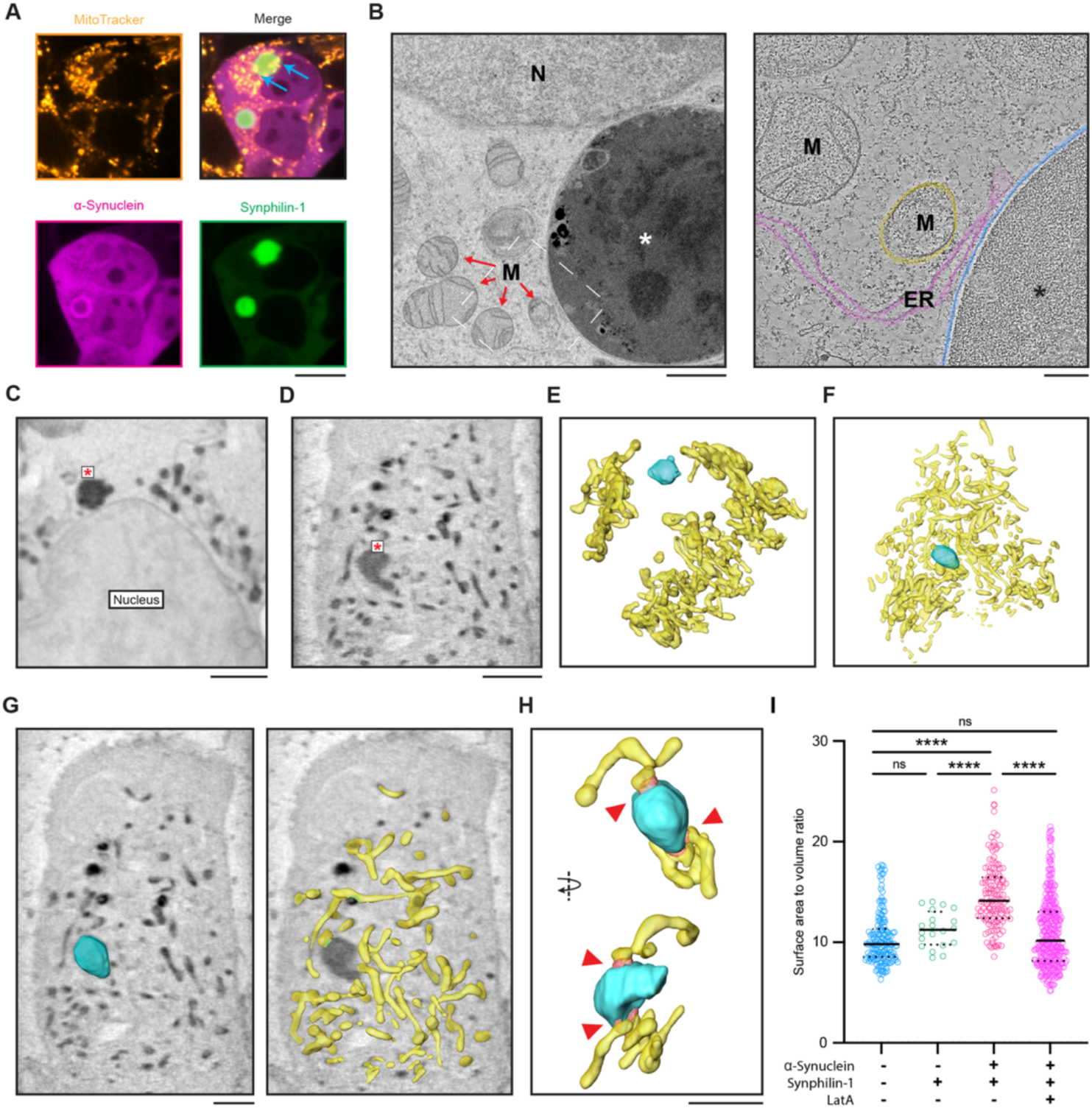
Lewy body-like condensates sequester and alter mitochondrial morphology at the interface of condensates in actin-dependent manner. **(A)** Live-cell confocal microscopy images of HEK293 cells expressing EGFP-synphilin-1 and α-synuclein-BFP, 24 h post-transfection and incubated with MitoTracker-647. Scale bar, 10 µm. **(B)** Representative electron micrograph of cryo-fixed HEK293 cells expressing LBLs (left) and magnified tomogram slice with its 3D rendered volume (right; magenta: ER, yellow: mitochondria, blue: LBL). Note accumulation of organelles at the condensate interface. Dashed line indicates region used for electron tomography. N indicates nucleus; M, mitochondria; ER, endoplasmic reticulum; *, LBL. Scale bar, 1 µm (micrograph), 200 nm (tomogram). **(C)** SXT orthoslice of a representative cell expressing EGFP-synphilin-1. Asterisk indicates synphilin-1 condensate. Scale bar, 2 µm. **(D)** SXT orthoslice of a representative cell expressing LBLs (EGFP-synphilin-1 and α-synuclein-BFP). Asterisk indicates an LBL. Scale bar, 2 µm. **(E)** Surface rendering of the cell in *C,* indicating the reconstructed mitochondrial network (yellow) around synphilin condensate (cyan). **(F)** Surface rendering of the cell in *D,* indicating the reconstructed mitochondrial network (yellow) around an LBL (cyan). Note the accumulation of mitochondria at the interface of an LBL. **(G)** SXT orthoslice of a representative tomogram of a cell expressing EGFP-synphilin-1 and α-synuclein-BFP demonstrates the accumulation of mitochondria at the interface of Lewy body-like condensates. Cyan, surface-rendered LBL; yellow, surface-rendered mitochondria network. Scale bar, 3 µm. **(H)** Magnified region indicates contacts between mitochondria and LBLs (red arrowheads). Scale bar, 3 µm. **(I)** Surface-to-volume ratios of mitochondria rendered from the SXT obtained from untransfected cells (blue), cells expressing synphilin-1 (green), LBLs-containing cells (magenta) and cells expressing synphilin-1 and α-synuclein and treated with Latrunculin A. Data are from three independent experiments. Significance was tested using one-way ANOVA test; ns, non-significant; asterisks, significance: ****, *p* < 0.0001.

One of the major strengths of SXT is the ability to reconstruct the isotropic 3D imaging of an intact cell without physically slicing it. This allowed us to determine the spatial relationship between organelles and intracellular structures up to 35 nm resolution in all three dimensions^52^. To validate SXT for condensate-to-membrane characterization, we turned to the well-described system of synaptic vesicle condensates by co-expressing synapsin-1 and synaptophysin. Here, SXT reliably recapitulated the vesicular cohorts specific for the ectopically generated SV-like condensates (**Supplementary Fig. 10A**). In line with the confocal imaging and classical electron microscopy^53^, the only accumulation of lipids was shown by FM 4-64 as small lipid vesicles colocalized with synapsin-1/synaptophysin condensates, mimicking the sequestering of vesicles at the synaptic boutons, while mitochondria formed stereotypical network in cells (**Supplementary Fig. 10B-10D**). These observations are in line with our live-cell imaging using organelle tracers, suggesting that correlation of light microscopy and SXT imaging datasets is a suitable approach to investigate the interfaces between membrane-bound organelles and biomolecular condensates in cells.

As our correlated measurements of light microscopy and soft X-ray tomography revealed that the maturation of LBLs occurs over time (**Supplementary Fig. 2C and 5C**), LBL condensates could function as a sink for actin and membranes sequestering. This motivated us to employ SXT to characterize the ultrastructure of mitochondria juxtaposed to LBLs. In line with results from light microscopy, we were able to observe a major difference in the mitochondrial network around the condensates (**Fig. 5C-5F; Supplementary Movie 3**). In the case of synphilin-only condensates, the mitochondrial network remained intact and similar to non-transfected cells. When expressing LBLs, the mitochondrial network sharply changed: (i) mitochondria appeared to be more fragmented in the vicinity of condensates (**Fig. 5G**), as indicated by an increased surface-to-volume ratio (**Fig. 5I**); (ii) the distance between the LBLs and mitochondria dropped and fragmented mitochondria accumulated at the surface of LBLs (**Fig. 5H; Supplementary Movie 4**). Inhibition of actin polymerization with LatA attenuated LBL formation and normalized the measured mitochondrial morphological parameter (**Fig. 5I**). Because LatA directly perturbs actin function and no cellular or neuronal functional endpoint was measured, this experiment should not be interpreted as demonstrating functional rescue.

Nevertheless, the demixing of α-synuclein appears to be associated with mitochondrial accumulation in this model, because synphilin-only condensates and unrelated synaptic vesicle-like condensates lack such accumulations. These observations demonstrate a direct relationship between α-synuclein demixing, actin organization, and mitochondrial morphology, irrespective of whether inclusions themselves being potentially injurious or the phenotype arising from depletion of soluble α-synuclein, actin, or other recruited components.

### Lewy body-like structures form in primary neurons and recapitulate molecular signatures of post-mortem LBs

Next, we asked whether LBLs form in primary neurons, where cellular context is substantially different. To address this, we co-transfected EGFP-synphilin-1 and α-synuclein-BFP in primary hippocampal neurons. We could observe the formation of early-stage mixed condensates and late-stage LBLs with the core-shell morphology (**Fig. 6A, 6B**) 24 h post-transfection. As with experiments in HEK293 cells, treatment of transfected neurons with LatA reduced the frequency of LBLs formation in neurons (**Fig. 6C**). We also performed staining against p62, both in neurons and HEK293 cells, which demonstrated the accumulation of p62 bodies on the surface of LBLs (**Supplementary Fig. 11**). These results demonstrate that LBLs are not restricted to non-neuronal cells. Instead, our results suggest that cells mark these LBLs for autophagic degradation.

**Fig. 6.**
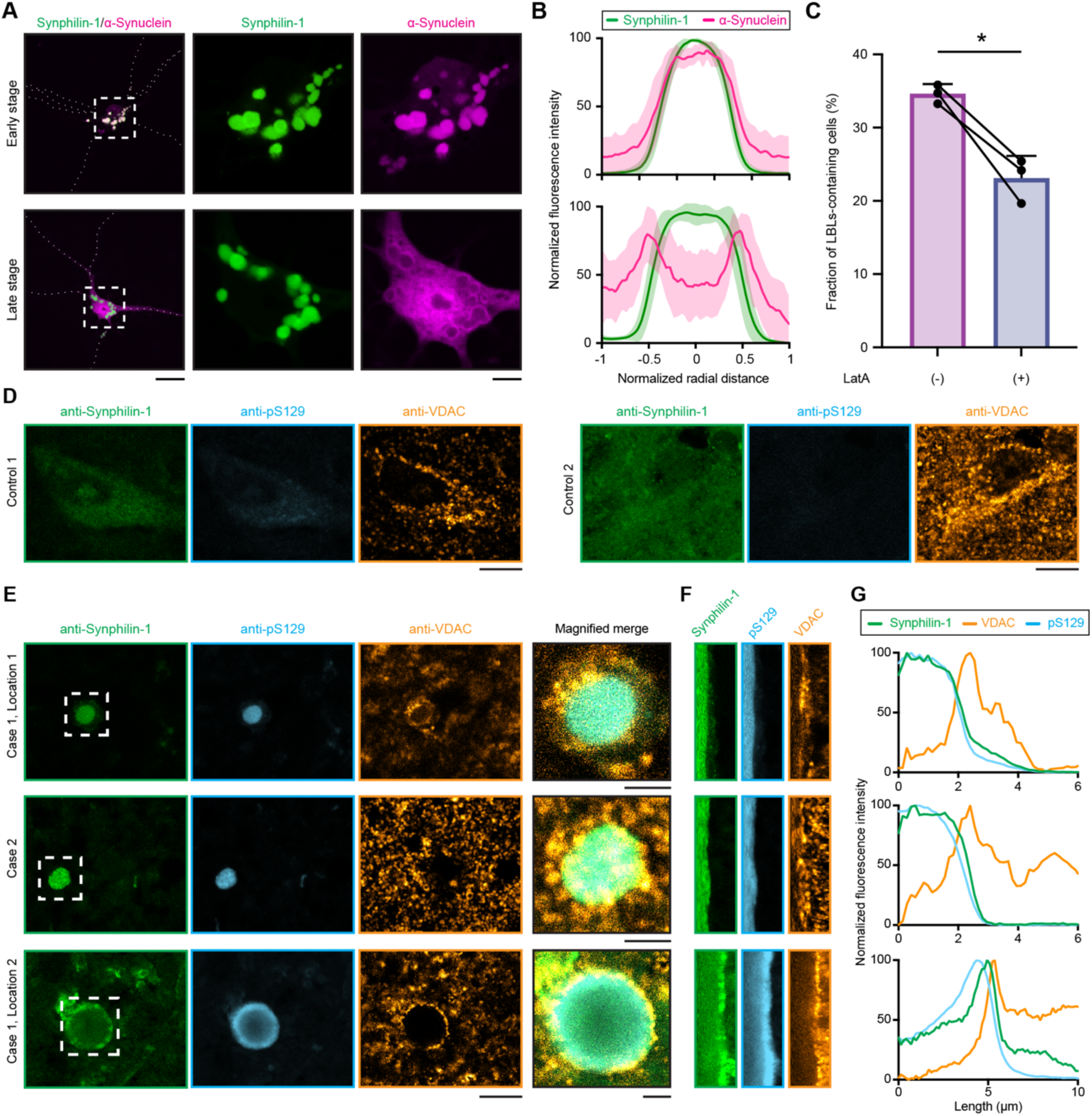
Lewy body-like condensates form in neurons and recapitulate selected structural features of human Lewy bodies. **(A)** Representative images of wild-type murine hippocampal neurons in culture expressing early-stage (top; EGFP-synphilin-1 (green) and α-synuclein-BFP (magenta) are fully mixed), and late-stage LBLs (bottom; EGFP-synphilin-1 and α-synuclein-BFP form core-shell morphology). White dashed squares delineate magnified regions; white dotted lines represent neurites. Scale bars, 20 µm (overview), 5 µm (magnified). **(B)** Cumulative line profiles for conditions in *A*. Line profiles were collected from 10 condensates across 3 biological replicates, each from a separate cell. Bold line represents mean value, shaded area represents SD. **(C)** LatA treatment inhibits LBLs formation in neurons, as represented by the fraction of LBL-containing cells. Data from at least 300 cells across three independent replicates for each condition. Error bars represent SD; dots represent individual replicates; lines connect values for DMSO control and LatA treatment from the same biological replicate. Significance was tested using a paired Student’s t-test; asterisks, significance: *, p<0.05. **(D)** Immunohistochemistry on paraffinized human brain tissues from individuals with no Lewy body pathology (control), stained against synphilin-1 (green), pS129 α-synuclein (blue), and VDAC channel on mitochondria (orange). Note the absence of LBs. Scale bars, 10 µm. **(E)** Representative examples of different stages of LB pathology in the substantia nigra and hippocampus from individuals with LB pathology stained against synphilin-1 (green), pS129 α-synuclein (blue), and VDAC channel on mitochondria (orange). Note the accumulation of mitochondria fragments at the interface of LBs in the merged images. In total, 11 LBs were analyzed from three individuals, 9 of which were positive for synphilin-1. Scale bars, 10 µm (overview), 3 µm (magnified). **(F)** Fluorescence intensity profiles across the interface of LBs shown in *E.* **(G)** Radial line profiles across the LBs from individual LBs shown in *E*, demonstrating the distribution density of mitochondria at the interface of LBs.

Synucleinopathies are increasingly perceived as “protein-induced lipidopathies”, suggesting that lipids and membrane-bound organelles may act as a trigger of α-synuclein aggregation^15^. In this scenario, the aberrant interactions between α-synuclein and membranes could occur early in the biogenesis of LBs, whereas amyloid fibrils are characteristic of a later stage with more mature inclusions. Several recent studies focused on the ultrastructure of α-synuclein-containing inclusions in patients, demonstrating that these inclusions are, in fact, a medley of membrane-bound organelles, such as mitochondria^10^. Yet these experiments do not investigate the mechanism by which mitochondria end up trapped at the interface of α-synuclein inclusions. We analyzed brain tissue from individuals with Lewy body pathology to test the hypothesis that the core-shell morphology of α-synuclein – synphilin-1 condensates relates to the accumulation of mitochondria. Staining of substantia nigra and hippocampus identified LBs at different stages, emphasizing the core-shell architecture of α-synuclein (**Fig. 6D, 6E**) and the accumulation of mitochondria at the periphery (**Fig. 6E-6G**). These findings confirm that the LBL system recapitulates the spatial accumulation of mitochondria at the periphery of inclusions observed in human brain tissue^16^. This end-point morphological correspondence does not establish temporal order, mechanism, or whether mitochondrial redistribution is pathogenic, compensatory, or secondary to another cellular disturbance. Nonetheless, these findings confirm that the LBLs recapitulate the sequestration of mitochondria, a hallmark of inclusions found in human brain tissues.

### Lewy body-like condensates are reaction centers for α-synuclein fibrillization

Quantitative spectroscopy analyses of LBs in post-mortem tissues of individuals with PD suggest that α-synuclein fibrils preferentially form at the surfaces of these pathological inclusions^54^. This aligns with the idea that condensates provide a unique chemical environment^55^, which can accelerate the aggregation of proteins, including α-synuclein^26,27,56,57^. Recent studies showed that the interfaces of condensates can establish electric potential gradients^58^, thereby encoding unique condensate microenvironments (e.g., redox state) that affect protein fibrillization^28,59^. To explore this possibility, we stained LBLs-expressing cells with Peroxy Orange 1, an activity-based sensor selective for H_2_O_2_. We observed increased fluorescence within condensates in comparison with surrounding cytoplasm. This observation indicates that Lewy body-like condensates maintain a redox environment distinct from the surrounding cytosol (**Supplementary Fig. 12**), which in turn may provide the chemical driving force for condensate-dependent fibrillization.

To further elucidate the beta-sheet content and aggregation state of the proteins within the LBLs, we employed scanning X-ray nanodiffraction analysis in LBLs-expressing cells (**Fig. 7A**). X-ray diffraction can be used to detect amyloid fibrils, which are characterized by an elongated cross-beta-sheets structure. This results in a sharp diffraction peak corresponding to the inter-strand spacing along the fibril axis (typically around *d* ≈ 4.7 Å), and a more diffuse peak corresponding to the inter-sheet spacing (typically around *d* ≈ 10 Å) perpendicular to the fibril axis^60^. Scan of dried α-synuclein preformed fibrils (PFF) produced a characteristic X-ray scattering profile with a sharp diffraction peak centered at *q* = 1.3544 Å^-1^(*d* ≈ 4.64 Å), expected from a highly ordered cross-beta structure (**Fig. 7B,7C**). This diffraction curve was therefore used as a reference for a fibrillization process.

**Fig. 7.**
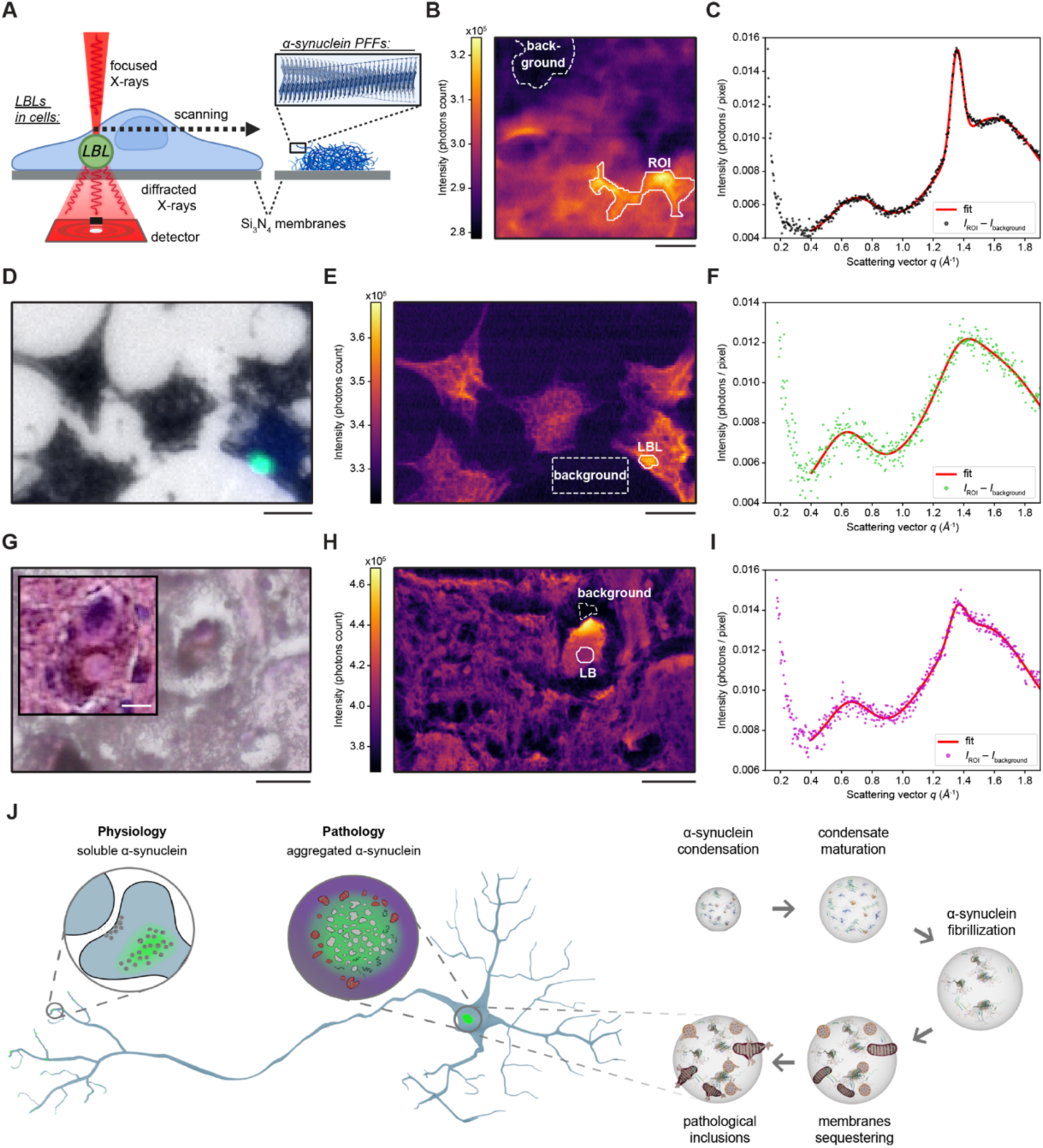
Scanning X-ray nanodiffraction analysis reveals fibrillization in Lewy body-like condensates. **(A)** Schematic illustrating the experiment setup. α-synuclein PFFs, HEK293 cells containing LBLs, and a histological section containing a LB from the substantia nigra of a PD patient were raster-scanned with a focused X-ray beam (beam diameter ≈ 250 nm, photon energy = 15 keV), recording a 2D scattering pattern in each scan point by a photon-counting pixel detector. **(B)** X-ray dark-field map of dried α-synuclein PFFs. The intensity was integrated between *q* = [0.4, 1.9] Å^-1^. Scale bar, 10 µm. **(C)** Average scattering profile of α-synuclein PFFs within the ROI selected in *B*. The sum of three Gaussians is fitted to the data. **(D)** Optical micrograph of freeze-dried HEK293 cells expressing EGFP-synphilin-1 and α-synuclein-BFP (composite image of the bright-field, EGFP, and BFP channels). An LBL is visible in the lower-right cell. Scale bar, 10 µm. **(E)** X-ray dark-field map of the sample from *D*. The intensity was integrated between *q* = [0.4, 1.9] Å^-1^. Scale bar, 10 µm. **(F)** Average scattering profile of the LBL from *E*. The sum of three Gaussians is fitted to the data. **(G)** Optical micrographs of a hematoxylin & eosin-stained histological section from the substantia nigra of a PD patient containing a LB. The section was air-dried immediately before the X-ray measurements, resulting in some cracks. The inset shows the same LB before drying. Scale bar, 25 µm (overview), 10 µm (inset). **(H)** X-ray dark-field map of the sample from *G,* showing the integrated intensity between q = [0.4, 1.9] Å-1. Scale bar, 25 µm **(I)** Average scattering profile of the LB shown in *H*. The sum of three Gaussians is fitted to the data. A description of the curve-fitting process and parameters of Gaussians for each curve are provided in the Methods section and Supplementary Tables S4-6. **(J)** Conceptual schematic summary of the study. Normally, α-synuclein is a soluble presynaptic protein with a physiological role. However, in pathology it is found in aggregates in neuronal soma and neurites. Our study provides a platform that faithfully recapitulates early stages of LB formation through phase separation, including fibrillization of α-synuclein and sequestration of membrane-bound organelles.

Correlative light microscopy/X-ray nanodiffraction analysis of LBLs expressed in HEK293 cells demonstrated a diffuse peak centered at *q* ≈ 0.62 Å^-1^ (*d* ≈ 10.1 Å), as well as a slightly asymmetric peak at *q* ≈ 1.37 Å^-1^ (*d* ≈ 4.6 Å) on top of the broader peak centered at *q* ≈ 1.55 Å^-1^ (*d* ≈ 4.1 Å) (**Fig. 7D-7F**). To assess whether this broadening was also observed in a more physiologically relevant context, we extended the analysis to patient-derived tissue samples. The diffraction signal of the LB from post-mortem human brain tissue (**Fig. 7G-7I**) exhibited a well-defined peak at *q* ≈ 1.358 Å^-1^, albeit with a smaller amplitude than that of the PFFs. This suggests the presence of some amyloid fibrils or higher-order aggregates, while the two broad peaks at *q* ≈ 0.65 Å^-1^ and *q* ≈ 1.51 Å^-1^ indicate the presence of other beta-sheet structures.

Although the amyloid diffraction peak in LBLs is not as sharp as in synthetic PFFs, it is comparable to the diffraction profile of patient Lewy bodies. Our data suggest the presence of beta-sheet structures in LBLs or at their interfaces, and the peak shoulder at *q* ≈ 1.37 Å^-1^ may appear due to shorter beta-sheet aggregates, indicating initial stages of the fibrillization process.

Soluble α-synuclein is a presynaptic protein with physiological functions, but its contribution to inclusion-associated cellular dysfunction remains unresolved. In the model described here (**Fig. 7J**), α-synuclein is recruited into condensates, co-condensing with scaffold proteins, undergoing progressive demixing while becoming less mobile. Actin and membrane-bound organelles accumulate within or around the resulting structures; β-sheet-rich assemblies arise subsequently, most likely at the interfaces. This trajectory identifies at least three nonexclusive potential sources of dysfunction: (i) depletion of physiologically available soluble α-synuclein or other recruited proteins; (ii) altered cellular organization associated with the inclusion; and (iii) later acquisition of fibrillar structures. The present study establishes a platform for distinguishing among these possibilities, without ranking their causal importance.

## DISCUSSION

We reconstituted LB-Like structures driven by co-phase separation of synphilin-1 and α-synuclein. These structures were reconstituted in vitro and in minimal cellular systems, and the key features have been validated in neurons and human post-mortem tissue. Proteomics, live imaging, and soft X-ray tomography show that these inclusions are associated with redistribution of actin and membrane-bound organelles and with altered mitochondrial morphology. Structural analyses further indicate that the condensates with core-shell organization can act as reaction centers for local nucleation of β-sheet structures, suggesting a pathway from pathological condensation to insoluble fibril generation.

### α-Synuclein condensation as a trajectory towards the formation of Lewy bodies

Functional condensates, such as the ones with synapsin-1 and synaptic vesicles, are essential for synaptic function^21,61–63^. Yet, the condensation driven by scaffold proteins, such as synphilin-1, outside the presynapse might lead to the formation of viscoelastic inclusions^64–66^. Such protein transitions have been reported for inclusions of hnRNPA1, FUS and TDP-43^67–70^. However, it is far less clear whether these different types of protein inclusions reflect the same pathological mechanism and require fibrils for its downstream effects. For example, α-synuclein-containing condensates may act as sinks of soluble α-synuclein molecules, such that recruitment, concentration, demixing, and progressive immobilization reduce the physiologically available α-synuclein pool. This possibility is particularly relevant because the study measures recruitment and reduced mobility but does not quantify the remaining soluble pool or its physiological activity. Thus, depletion is a mechanistically plausible interpretation rather than a demonstrated outcome. In our reconstitution system, we find that α-synuclein progressively demixes, forming a characteristic core-shell structures that are highly reminiscent of Lewy bodies found in patient tissues^10,11,16^. This demixing occurs only when a threshold concentration of synphilin-1 is reached in the condensates (**Fig. 1**), resulting in a stable shell of low solubility that re-mixes only under extreme conditions such as in a hypotonic solution (**Supplementary Fig. 6**). Such concentration-dependent demixing is a characteristic of phase-separation ^71,72^. Together, our findings suggest a trajectory from phase separation of α-synuclein towards the formation of Lewy bodies and that inclusion formation can be mechanistically separated from fibrillization, arguing against treating fibril generation as the necessary primary pathogenic event.

### Intracellular organelle trafficking at the crossroad of Lewy body-like assemblies

Unbiased proteomics analysis of LBLs-expressing cells demonstrated broad changes in protein landscape with particular downregulation of cytoskeleton-related proteins. Live-cell imaging with fluorescent trackers further showed that actin, but not tubulin, was sequestered within LBLs (**Fig. 4**). Both correlative light and electron microscopy and soft X-ray tomography showed a striking accumulation of fragmented mitochondria at the surface of LBLs (**Fig. 5**), consistent with earlier CLEM studies of patient Lewy bodies^16^ and our own histopathology data from patient tissues (**Fig. 6**). This all-or-none mitochondrial enrichment around aberrant inclusions – a feature observed in several neurodegenerative diseases^73^ – as well as assembly of LBLs themselves, was dependent on actin sequestration and was accompanied by disruption of mitochondrial network morphology. One potential explanation could be that local collapse of the actin network perturbs mitochondrial dynamics, altering the balance between mitochondrial fission and fusion and disrupting short-range mitochondria trafficking, which leads to their sequestration at the surface of LBLs. Such disruptive effects on mitochondrial network depend on α-synuclein, as synphilin-only condensate or unrelated synaptic vesicle-like condensates lack these organelle accumulations.

Finally, the accumulation of p62-positive bodies at the surface of LBLs indicates that they are targeted for autophagic degradation, giving rise to a potential mechanism where p62 recruitment and organelle sequestration represent cellular containment or disposal responses. These features should not automatically be interpreted as evidence that the inclusion is the initiating toxic species. The actin sequestering and mitochondrial fragmentation alone cannot establish whether the relevant injury comes from the inclusion itself, depletion of soluble α-synuclein, or both. However, the fact that LB-like system described here recapitulates the organelle redistribution and mitochondrial fragmentation allows for designing functional rescue experiments. One step and a proof-of-principle for such rescue approach has been the acute inhibition of actin polymerization, which attenuated the formation of LB-like structures in both ectopic system and in neurons (**Figs. 4F and 6A-6C**), rescuing the mitochondrial network (**Fig. 5I**).

More broadly, the entire condensation process may constitute a stress-induced containment response to an antecedent cellular insult. In that scenario, p62 recruitment, organelle redistribution, and protein immobilization would be components or consequences of the response rather than evidence that the inclusion is the original vehicle of disease.

### Implication of α-synuclein-containing condensates for the formation of insoluble cross β-sheet fibrils

The relationship between condensates and the formation of beta-fibrils remains debated: for some types of condensate-forming proteins formation of fibrils has been suggested as an off-path reaction, whereas for others the condensates are thought to be necessary for the fibrillization, either within the condensates or at the interface of condensate and surrounding phase^74–77^. Data from Peroxy Orange 1 reporter suggest that LB-like inclusions contain higher redox potential, which can catalyze fibrillization^78^. Scanning fluorescence imaging/x-ray diffraction studies support the notion of LBs and LB-like inclusions harnessing beta-sheet structures (**Fig. 7**). Even in a complex human tissue with LBs, the diffraction signal contained a peak reminiscent to the peak seen in LB-like inclusions at the related position where a sharp peak typically associated with pure amyloid fibrils would be. The proposed sequence – from soluble protein to condensation, maturation, and only subsequently beta-sheet formation – is consistent with insoluble pathology being the endpoint of depletion from the soluble pool. Moreover, scanning X-ray diffraction studies of human tissue LBs^79^ found that only a subset of LBs exhibited a diffraction pattern characteristic of amyloid fibrils, while others lacked definitive features of cross-beta structures. Together with broad histopathological definition of Lewy body itself^80^, these observations suggest that Lewy bodies are structurally heterogeneous and may contain a spectrum of protein conformations, ranging from assemblies enriched in cross-beta structure to those with little or no detectable cross-beta organization. The protein organization observed in our LB-like system may enable detailed investigation of LBs, organelle trafficking and the disease phenotype irrespectively of the seeding process or fibril formation.

The observation that the condensation leads to the generation of α-synuclein-containing inclusions, which sequester actin, may deplete the soluble α-synuclein pool and are associated with mitochondria disruption, illustrates the critical role condensates can have in regulating intracellular membrane trafficking (**Fig. 7**). These relationships do not establish that the condensate initiates toxicity. Condensation may instead impair cellular physiology by withdrawing α-synuclein, actin, and other proteins from their soluble functional pools. Conversely, recruitment of organelles and p62 may represent containment or disposal responses. A future experiment may be dedicated to measuring soluble versus condensate-associated α-synuclein over time and determine whether replenishing the functional soluble pool improves cellular physiology even when inclusions remain. That would directly discriminate between the gain-of-inclusion and loss-of-soluble-function possibilities. While LBs contain various proteins^11,80,81^, the minimal system of α-synuclein and synphilin-1 described here provides a long-sought-after cellular proxy for LBs. Such a system, in conjunction with a large-scale compound screen, can identify small molecules that are able to modulate viscoelastic properties of condensates^82^, allowing for the development of new therapeutic strategies. In a broader scope, functional coupling of membrane-bound organelles and biomolecular condensates is an emerging general property for the intracellular organization^83,84^.

### Limitations of the Study

While the minimal system of LBL inclusions captures the key features of patient, several limitations need to be acknowledged. It is important to note that, while the human findings establish morphological correspondence but not temporal order or causation, the postmortem similarity cannot determine whether Lewy bodies caused neuronal dysfunction, arose in response to it, or became harmful by progressively depleting soluble proteins. Additionally, although our LB-like structures have the key morphological features of LBs, the use of synphilin-1 as a driver does not capture the possible influences from additional binding proteins that may critically contribute to functional outcomes by being trapped in these condensates and depleted away from their functional, soluble pools. The upstream cause of α-synuclein condensation also remains unresolved. In human disease, condensation may be a reactive phase transition triggered by toxic, metabolic, inflammatory, infectious, or other biological exposures that disturb proteostasis or the cellular environment. Thus, the ability of condensation to generate LBL structures does not establish condensation as the initiating disease mechanism. The resulting inclusion could be protective, maladaptive, or initially protective but ultimately consequential through progressive depletion of soluble functional proteins. Lastly, this study did not address whether the selective vulnerability of specific dopaminergic neuron subtypes is directly correlated with LB/LBL formation. Future work using iPSC-derived, subtype-specific dopaminergic neurons will be required to systematically investigate this relationship.

## METHODS

### DNA construct preparations

#### Primer design

Primers were purchased from Sigma Aldrich, dissolved in ddH2O (100 µM stock concentration) and stored at −20 °C. All primers used in this study are shown in the supplementary table 2.

#### Synphilin-1 plasmids

Plasmid for EGFP-synphilin-1 contained the human sequence for synphilin-1 CDS with an N-terminal EGFP tag in a standard pEGFP-C1 backbone (Supplementary table 1). For protein purification of synphilin-1 in mammalian cells (Expi293F^TM^), the sequence of EGFP from the synphilin-1 plasmid was replaced using restriction enzyme sub-cloning with *Age*I-*Xho*I by an expression cassette consisting of a 14X Histidine affinity, SUMO_Eu1- and EGFP cassette tag. The His-14-SUMO_Eu1 tag was taken from pAV0279 (Addgene 149688) (Supplementary table 1)^85^.

#### α-Synuclein constructs for recombinant purification

For α-synuclein purification from bacteria, WT and 140C α-synuclein were expressed from pET28a plasmid. The pET28a expression vector was created by subcloning α-synuclein (Addgene #51437) with restriction enzyme sites *Nco*I-*Not*I into the pET28a vector. For subsequent fluorescent labeling, alanine 140 was replaced with cysteine by mutagenesis PCR (A140C) according to standard mutagenesis PCR method using primers #DML0463 & 0464.

#### α-Synuclein mutants for live-cell imaging

To obtain C-terminally tagged α-synuclein-BFP, α-synuclein (Addgene #51437) was PCR sub-cloned into a BFP-N1 vector (kind gift from Geert van den Bogaart, Groningen, NL) using restriction sites *Bam*HI-*Xho*I^21^. To obtain the truncated variants, α-synuclein ΔF and α-synuclein ΔC, single-fragment Gibson Assembly cloning was used. In brief, truncated Δ-α-synuclein-BFP was amplified from template pBFP-N1 SNCA by PCR with primers #DML0486 & 0487 (ΔF), or #DML0492 & 0493 (ΔC) respectively. The methylated parent template plasmid was digested with 10 U *Dpn*I for at least 1 hour at 37 °C. Reactions were analyzed on preparative 1% agarose gel electrophoresis. Subsequently, 50 ng of the gel-purified PCR product was used in Gibson Assembly reaction using an in-house mix (0.03 U/µL Phusion® 0.005 U/µL T5-exonuclease, 5.3 U/µL Taq-DNA ligase in 100 mM Tris-HCl pH 7.5, 5 mM MgCl_2_, 0.2 mM dNTP each, 10 mM DTT, 2.5% (w/v) PEG 8000 and 1 mM NAD^+^) incubated for 1 hour at 50 °C. Reactions were directly used for the transformation of chemically competent bacteria. All the vectors contain CMV promoter and SV40 terminator.

The PD-associated α-synuclein point mutations (A30P, S129E) were introduced in pBFP-N1 SNCA plasmid by standard mutagenesis PCR. Primer pairs were designed in an overlapping fashion covering the point mutation region of interest: 10 bp upstream and at least 17 bp downstream of point mutation site were complementary to the plasmid template. All plasmid sequences were verified by Sanger DNA sequencing.

#### α-synuclein-Halo construct for AAVS1 CRISPR-Cas9 integration

α-synuclein-Halo fragment and SV40 terminator sequence fragment were amplified from existing α-synuclein-Halo vector (kind gift from Helge Ewers, Berlin, DE) with primers #DML562 & #DML563 and #DML564 & #DML565 respectively. Target plasmid vector (amplified from modified AAVS1 integration vector (Addgene #92099), containing additional multiple cloning site) was digested with 10U of BcuI restriction enzyme in Buffer O at 37 °C for 1 hour. Final fragments were assembled via Gibson Assembly. Assembled vector was transformed in DH10B E. coli cells, plasmids from several clones were purified and sent for sequencing. Plasmid with correct sequence was then used for gene integration in cell line.

*Additional Constructs.* For control experiments, we used tag-free synaptophysin^53^.

### Cell culture

#### Transfection of HEK293 cells using Lipofectamine 2000

HEK293 cells were transfected by Lipofectamine 2000 (Thermo Fisher) following the manufacturer’s instructions. Briefly, 3 µl of Lipofectamine 2000 was mixed with 2 µg of total DNA in 200 µl OptiMEM (Gibco). The transfection mix was incubated for 30 min at room temperature and added to the cells. Cells were transfected and incubated overnight (37 °C and 5% CO2). The medium was fully replaced with fresh-supplemented DMEM one day post-transfection.

#### Immunohistochemistry in cells

Cells were fixed 18 or 22 h after transfection with 4% PFA (Thermo Fisher 28908) in PBS (Gibco 10010015) for 15 min at 4°C, permeabilized in buffer consisting of 0,1% saponin (Sigma 47036), 3% BSA (Sigma A7906) and 10 mM glycine (Roth 2908) for 30 min at 4°C and stained with anti-pS129 α-synuclein (D1R1R, Cell Signaling) or anti-p62 (ab109012, Abcam) primary antibodies. Dilutions used are 1:500 for anti-pS129 and 1:400 for anti-p62. Cells were incubated with antibodies overnight at 4°C and washed with 0,1% saponin in PBS. Following this, cells were stained with secondary Alexa Fluoro 568 anti-rabbit antibodies (A11011, Invitrogen) at room temperature for 45 min, washed with 0.1% Saponin in PBS and mounted with Fluoroshield (Sigma F6182) on a glass slide for imaging. Based on live-cell longitudinal imaging, “early stage” category was assigned to LBLs with mixed α-synuclein and synphilin-1 signal; “late stage” – to LBLs with demixed α-synuclein.

#### Preparation of hippocampal neurons

Hippocampal neurons were prepared from P0/P1 wild-type, C57BL6/J mouse line. Brains were dissected out and hippocampi were collected in cold Hank’s balanced salt solution (HBSS, Gibco) supplemented with 10 mM HEPES (Gibco), 1 mM Pyruvic Acid (Gibco), 0.5% Penicillin/Streptomycin and 5.8 mM MgCl_2_. Upon dissection, hippocampi were enzymatically digested with papaine (Sigma) in HBSS for 20 min at 37 °C. Subsequently, papaine was exchanged by the plating medium (Neurobasal Medium A (Gibco, A2477501), supplemented with 5% FBS, 1% B27, 1x Glutamax, and 1% Penicillin/Streptomycin). The final cell suspension was obtained by mechanical dissociation with a P1000 pipette. Cells were seeded on glass coverslips pre-coated with 0.1 mg/mL poly-L-lysine (PLL; Sigma). Neurons were maintained at 37 °C and 5% CO2 in Neuronal Media (NB-A supplemented with 1% B27, 1% Glutamax, and 0.5% penicillin/streptomycin) and half-media changed every two days. Live-cell imaging experiments were performed on neurons that were at least 14 days in culture. All animal experiments were approved by the Institutional Animal Welfare Committees of the Charité University Clinic (Berlin, DE).

#### Transfection of neurons using Lipofectamine 2000

Neurons were transfected with Lipofectamine 2000 (Invitrogen, 11668500) following the modified manufacturer’s instructions. Coverslips with neurons were first transferred in a new dish with NB-A medium (Gibco, A2477501). For each transfection 3 μl of Lipofectamine 2000 was mixed with 1 μg (single transfections) or 2 μg (double and triple transfections) of total DNA in 200 μl of NB-A. The transfection mix was incubated for 20 min at room temperature and added to the cells. After 1 h coverslips were returned to original dishes and neurons were maintained in a humidified incubator at 37 °C and 5% CO2 for 24 hours (if not indicated otherwise) before following procedures.

### Stable cell line generation

#### CRISPR-Cas9-mediated knock-in and generation of α-synuclein-Halo stable cell line

Stable α-synuclein-Halo cell line was generated by CRISPR-Cas9-mediated integration into the AAVS1 safe-harbor locus. For this purpose, cells were transfected using Lipofectamine 2000 with a combination of (i) an AAVS1 donor integration vector containing the α-synuclein-Halo cassette and (ii) a complementary plasmid encoding the guide RNA template together with Cas9. Twenty-four hours after transfection, the medium was replaced with fresh DMEMc supplemented with 2 µg/mL puromycin (Thermo Scientific, A1113803), which was maintained throughout the whole subsequent selection procedure. After 72 hours of selection, cells were seeded into 96-well plates by limiting dilution at a density of 0-1 cell per well. Wells were inspected after 24 h, and those containing a single cell were identified and used for clonal expansion. Clones were maintained in 96-well plates until sufficient growth was achieved, followed by stepwise expansion into 24-well and 6-well plates. For monoclonality screening, clones in 6-well plates were reseeded into replicate plates and induced 24 hours later with 500 ng/mL doxycycline. After an additional 24 h, cells were labeled with JaneliaFluor JF549 HaloTag ligand (Promega, HT1020, 1:1000) and imaged using an inverted Nikon Eclipse Ti2 microscope equipped with a Nikon Ph1 ADL 10×/0.25 NA air objective and an ImagingSource DMK 33UX174 CMOS camera. Clones exhibiting uniform fluorescence without detectable subpopulations of non-expressing cells were further expanded in T25 and T75 flasks.

Selected clones were seeded in 6-well plates and induced with 500 ng/mL doxycycline 24 hours after seeding. 24 hours after induction lysates were collected in RIPA buffer (Sigma, R0278). In parallel, HEK293 cells were seeded on 25mm glass coverslips. 24 hours later cells were transfected with Halo-synapsin-1 and α-synuclein-Halo using Lipofectamine 2000. Cells were stained with CellMask Actin Green (Invitrogen, A57243) and JaneliaFluor JF549 HaloTag ligand (Promega, HT1020, 1:1000) 24 hours post-transfection and imaged on Nikon spinning-disk confocal CSU-X (SDC CSU-X) microscope equipped with two EMCCD cameras (iXon3 DU-888 Ultra), Andor Revolution SD System (CSU-X), and a Plan Apo163 60x/1.4 NA, oil immersion objective. Immediately after imaging cells lysates were collected in RIPA buffer. Transfection efficiency was then calculated as the number of fluorescent (expressing) cells divided by the total cell number (determined by actin staining).

Protein expression in stable clones was assessed by Western blotting for the integrated protein (synapsin-1 or α-synuclein) and actin, in parallel with lysates from transiently transfected HEK293 cells. Band intensities were quantified using the built-in GelAnalyzer plugin in FIJI, normalized to actin, and corrected for transfection efficiency. The clone exhibiting the highest normalized expression level was selected as the final stable cell line.

#### Western blot analysis of stable cell line

Lysate samples were separated on 12% SDS-PAGE gels and transferred to 0.2 µm nitrocellulose membranes (Cytiva, 10600001) by semi-dry transfer at 200 mA at RT for 2 hours. Membranes were cut according to molecular weight prior to blocking and antibody incubation. Membranes were washed in TBS and blocked for 1 hour at room temperature in either 5% (w/v) milk (BD, 232100) in TBS. Primary antibodies were incubated overnight at 4°C with gentle agitation. After washes in TBS-T and TBS, membranes were incubated with species-appropriate HRP-conjugated secondary antibodies. Primary antibodies used: anti-SDHA (Millipore, MABN630, 1:1000 in 5% milk/TBS) and anti-α-synuclein (Abcam, ab53726, 1:500 in 5% milk/TBS). HRP-conjugated secondary antibodies used: goat anti-mouse (Sigma, A4416, 1:5000) and goat anti-rabbit (Sigma, A6154, 1:5000). Blots were developed using Amersham ECL Prime Western Blotting Detection Reagents (Cytiva, RPN2232) and imaged on Fusion FX device (Vilber).

#### Junction PCR

Junction PCR validates the integration of a cassette in a targeted genomic site. To this end, PCR was performed at 5’ and 3’ end of integration cassette, with one primer annealing within the integrated part and the other annealing to the genome outside of homology arm. Target sequences were amplified from plasmid templates using Phusion High-Fidelity DNA Polymerase and primer pairs #DML0615 & #DML0616 and #DML0614 & #DML0709 for 5’ and 3’ junctions respectively. 100ng of template gDNA were used. After reaction setup, samples were subjected to thermocycling using the touchdown program. PCR products were subsequently analyzed by agarose gel electrophoresis.

#### AK-11 insert sequencing

To validate the sequence of α-synuclein-Halo integrated in AK-11 genome, it was amplified from the genomic DNA using Phusion High-Fidelity DNA Polymerase and #DML0712 & #DML0713 primers. 100ng of template gDNA were used. After reaction setup, samples were subjected to thermocycling using the touchdown program. 5 µL of the PCR product were subjected to analytical gel electrophoresis and after confirmation of successful PCR the rest was digested with 10U of EcoRI restriction enzyme (Thermo Scientific, ER0271) in EcoR buffer (supplied with enzyme) at 37 °C for 1 hour. In parallel, the same digestion was performed on unrelated vector which does not contain neither α-synuclein nor Halo-tag sequence. After digestion vector was additionally dephosphorylated by 10 min incubation at 37 °C with 1U of FastAP (Thermo Scientific, EF0651) added to the reaction mix. Digestion products were run in 1% agarose gel and bands of corresponding sizes were cut out with subsequent DNA extraction. 50ng of purified vector fragment and 3x molar excess of purified α-synuclein-Halo insert fragment were ligated by incubation with 5 U of T4 Ligase (Thermo Scientific, EL0011) at 37 °C for 1 hour. Reaction mix was then transformed in DH10B E. coli cells, plasmids from several clones purified and sent for sequencing.

### Protein expression and purification

#### EGFP-synphilin-1

His-SUMO-EGFP-synphilin-1 was expressed in Expi293F^TM^ cells (Thermo Fisher Scientific) from the pCMV vector for 3 days following enhancement. Cells were harvested and lysed in a buffer that contained 25 mM Tris–HCl (pH 7.4), 300 mM NaCl, 0.5 mM TCEP (buffer A), and protease inhibitor (Complete EDTA-free, Roche). All purification steps were carried out at 4 °C. The lysates were centrifuged for 1 hour at 20,000xg, followed by a two-step purification. The first step was affinity purification on a Ni-NTA column (HisTrap™ HP, Cytiva) using 25 mM imidazole for the binding of proteins, 40 mM during wash steps, and 400 mM imidazole for elution of proteins (all in buffer A). Eluates were concentrated and subjected to size exclusion chromatography (Superdex™ 200 Increase 10/300, Cytiva) in 25 mM Tris-HCl (pH 7.4), 300 mM NaCl, and 0.5 mM TCEP (buffer B). The His-SUMO tag was cleaved using SUMO proteases (SENP1^EuB^) (1:100) overnight at 4 °C and the tag was removed by reverse Ni-NTA chromatography using cOmplete™ His-Tag purification resin (5893682001, Roche). The SUMO protease was recombinantly purified as described previously (pAV0286; Addgene 149333)^85^. Additionally, the affinity purified SUMO protease was subjected to a final polishing step using size exclusion chromatography (HiLoad™ 16/600 Superdex™ 200 pg, Cytiva) in 45 mM Tris-HCl (pH 7.4), 290 mM NaCl, 4.5 mM MgCl_2_, 2 mM DTT.

#### α-Synuclein WT and A140C

Untagged human α-synuclein and α-synuclein-A140C variant were expressed from pET28a vector (Novagen). In brief, a 1 L expression culture (LB; 10 g/L tryptone, 5 g/L yeast extract, 5 g/L NaCl) containing 50 µg/mL kanamycin was inoculated with an overnight preculture to achieve a final OD_600_ of 0.1. Cells were incubated at 37°C under shaking conditions and protein expression was induced at an OD_600_ of 0.7-0.8 with 1 mM isopropylthio-β-galactoside (IPTG). Cells were shifted to 18°C and incubated overnight (at least 16 h) under shaking conditions. After expression, cells were harvested by centrifugation at 7,000xg for 15 min (4°C). For α-synuclein purification the pellet was resuspended in 20 mL ice-cold PBS (Gibco 10010015) supplemented with protease inhibitors (Complete EDTA-free, Roche) and a spatula tip of lysozyme (A3711, AppliChem). Suspension was subjected to lysis in a French press. The lysate was cleared by centrifugation for 30 min at 50,000xg (Beckman Coulter Avanti J-E with rotor JA25.50) at 4°C. Nucleic acids were precipitated by adding streptomycin sulfate (S6501, Sigma) to 10 mg/mL final concentration with constant agitation for 30 min at room temperature.

For purification of α-synuclein-A140C variant 1 mM DTT was added. Subsequently the nucleic acids were pelleted by centrifugation at 50,000xg for 30 min (4°C). The supernatant was transferred to a clean glass beaker and proteins were precipitated by slowly adding ammonium sulfate (3746.1, Carl Roth) to a final concentration of 0.36 g/mL with constant agitation for 60 min at 4°C. Proteins were collected by centrifugation for 30 min at 50,000xg (4°C). The pellet was resuspended in 10 mL PBS and boiled for 20 min with stirring every few min. After centrifugation (30 min at 50,000xg, 4°C), the α-synuclein containing supernatant was subjected for dialysis against 5 L of 15 mM Tris-base solution overnight (132655T, Spectrum™ Labs Spectra/Por™1 6-8kDa MWCO Standard RC). For dialysis and subsequent ion exchange chromatography of α-synuclein-A140C variant 1 mM DTT (final concentration) was used in all steps. The dialyzed protein solution was passed through a 0.22 µm filter (10462200, Whatman® Puradisc 0.2 µm, Sigma) and subjected to anion exchange chromatography (HiPrep Q FF 16/10, Cytiva) in 25 mM Tris-HCl pH 7.7 (Buffer IEX). Elution was performed stepwise against 25 mM Tris-HCl pH 7.7 containing 500 mM NaCl (IEX-HIGH). First unwanted material was washed out with 30% IEX-HIGH, second α-synuclein was eluted with 50% IEX-HIGH (= 250 mM NaCl). Elution fractions were analyzed by SDS-PAGE (18% gel). Fractions containing α-synuclein were concentrated (Amicon® Ultra 4 MWCo 3kD, Merck Millipore) and subjected to size exclusion chromatography (Superdex™ 200 Increase 10/300, Cytiva) in 25 mM Tris-HCl (pH 7.4), 150 mM NaCl, 0.5 mM TCEP. α-Synuclein concentration was determined at 280 nm by using a molar extinction coefficient of 5960 M^-1^ cm^-1^. Proteins were snap-frozen in liquid nitrogen and stored at −80 C° until further use.

#### Labeling of proteins

For fluorescence labeling of α-synuclein, α-synuclein-A140C was thiol-maleimide conjugated with fluorescent Alexa 568 and Alexa 647 dyes from Thermo Scientific (Alexa Fluor™ 568 C5 Maleimide and Alexa Fluor™ 647 C2 Maleimide, 10 mg/ml). The labeling reaction was carried out in a buffer containing 150 mM NaCl, 25 mM Tris-HCl pH 7.2, and 1.3 mM TCEP for 2 h at RT with a 3 times molar excess of Maleimide dye. The unconjugated remaining dye was removed by running the labeling reaction over a PD-10 desalting column with Sephadex G-25 resin and elution with PBS (Sephadex™G-25 M, Cytiva). The degree of protein labeled was determined using the molar extinction coefficient and maximum absorbance length of respective dyes.

### Analysis of human brain samples

#### Postmortem human brain tissue cohort

Substantia nigra and hippocampus from patients with Lewy body pathology in the setting of Dementia with Lewy bodies or Alzheimer Disease and non-diseased control individuals (Table S3) were provided by the Center for Human Brain Discovery brain tissue biorepository at Yale University School of Medicine, Pathology Department. Informed consent for autopsy and research on post-mortem brain tissue was obtained by the next of kin in accordance with local and federal laws and procedures reviewed by Yale University’s Institutional Review Board. Study participants were allocated into disease or control groups based on clinical diagnosis and brain pathology as previously described^87^.

#### Immunohistochemistry on paraffinized human brain postmortem tissues

Tissue sections were deparaffinized with xylene (3 x 5 min each) and rehydrated in descending alcohol grades of 100%, 95%, 70%, and 50% for 10 min each before rinsing with running tap water and distilled water for 5 min. Air-dried slides were immersed in 1X citrate buffer at pH 6.0 in a pressure cooker with high steam for 4 min followed by natural pressure release (20 min) and then air-dried. Tissue was permeabilized in 0.2% PBST at room temperature for 15 min, followed by blocking with 3% BSA in 0.2% PBST for 30 min. The sections were then incubated with a primary antibody cocktail of rabbit anti-Synphilin-1 (Thermo Fisher BS-1905R, 1:200) and mouse anti-VDAC (Abcam, ab14734, 1:150) prepared in blocking solution at 4°C for 24 h. Sections were washed with PBS (3x), and incubated with anti-mouse Alexa Fluor™ 647 (1:200) (Invitrogen, A32728) and anti-rabbit Alexa Fluor™ 488 (1:200) (Invitrogen, A11034) at room temperature for 2 h and subsequently with Alexa Fluor® 594 anti-α-synuclein (phospho S129 antibody [EP1536Y] Abcam, AB282652, 1:50). After three additional PBS washes, sections were treated with autofluorescence quencher (Sigma cat no. 2160) for 5 min, and washed in 70% ethanol. Air-dried slides were mounted in Prolong Diamond antifade media and cured at room temperature for 24 h.

Three cases of Lewy body pathology and two control cases were analyzed. Images were captured from 10 fields for each case. Among these, approximately 4 fields showcasing structures exhibiting positive staining for anti-α-synuclein (pS129) were carefully selected. These structures displayed intracytoplasmic localization, with adjacent nuclear DAPI staining and a diameter of 5-30 μm, thereby meeting the morphological criteria characteristic of Lewy bodies^81,88^. Subsequently, 9 out of the 11 Lewy bodies identified were found to stain positive for synphilin-1. The detailed overview is provided in Supplementary Table 3.

### Imaging

#### Spinning disk confocal

Spinning disk confocal (SDC) microscopy was carried out on an Eclipse Ti Nikon Spinning Disk Confocal CSU-X, equipped with OkoLab Live-cells incubator (for control of temperature at 37 °C, 5% CO2), 2 EM-CCD cameras (AndorR iXon 888-U3 ultra EM-CCD), Andor Revolution SD System (CSU-X), objectives PL APO 60 /1.4NA oil immersion lens. Excitation wavelengths were: 405-nm for BFP; 488-nm for EGFP; 561-nm for Halo7(JF549), Alexa Fluor™ 568, FM 4-64, SPY555-Actin and mCherry; 647-nm for Alexa Fluor™ 647, LysoTracker™ Deep Red, MitoTracker™ Deep Red FM, SiR-actin and SiR-tubulin. Glass coverslips (1.5 thickness, 25 mm diameter) for live-cell imaging, glass-bottom dishes (1.5 thickness, 35mm diameter) for long-term live-cell imaging or 15-well 3D glass bottom slides (Ibidi) for in vitro samples were placed in a microscopy adaptor and then mounted on the objective stage. Exposure time 200 ms, gain 300 (EM gain 20 mHz 16-bit) and Piezo stage z-motor were used to collect z-series. Images were acquired using Acquisition software NIS Elements 5.21.02 and then analyzed with ImageJ (NIH). For circularity analysis, a mask of synphilin-1 condensates was created from a z-projection by thresholding. Particles were analyzed and divided into groups by their circularity (≤0. 89 or ≥0.9-1.0). For cumulative line profile analysis, the midpoint of a fixed line was aligned to the center-point of the condensate (EGFP-synphilin-1 signal), in order to align the line profile to each other.

#### FRAP in cells

Fluorescence recovery after photobleaching (FRAP) experiments were performed in HEK293 cells expressing α-synuclein-BFP and EGFP-synphilin-1, or EGFP-synphilin-1 alone. The bleach ROI was fixed to 1.12 µm of diameter and bleached at 100% transmission intensity (405 nm = 3.5 mW and 488 nm = 3.8 mW) for 500 ms and was positioned in correspondence on the shell of condensate (analyzed ROI diameter: 0.5 µm). For LBLs specifically where the two proteins, α-synuclein and synphilin-1 overlap (if not indicated differently). Reference ROI were defined in the cytosol of the same cell, and the Background ROI were defined outside of the cell. Recordings were carried out starting with 5 images pre-bleach, followed by a bleaching event and a post-bleach timelapse series of images (1 s frame rate) up to 60 s. Each FRAP experiment was performed in at least three independent biological replicates. Intensity recovery traces were obtained from the regions of interest were background corrected and all traces were normalized. All data are presented as mean ± s.e.m.

#### Long-term imaging of LBLs formation

HEK293 cells expressing EGFP-synphilin-1 and α-synuclein-BFP were grown in 35-mm glass bottom dishes. 12 h after transfection cell medium was replaced with fresh supplemented DMEM and dishes were mounted on the microscope stage. Images were acquired every 15 min for 12 h. Images of cells that did not contain condensates at the beginning, but formed them during imaging, were used for further analysis. Images from three independent transfections were collected.

### In vitro reconstitution assays

#### Reconstitution of synphilin-1 and phase diagram

For phase separation experiment, purified EGFP-synphilin-1 was mixed in different concentrations (1 µM, 5 µM, 10 µM), and different percentages of PEG 8000 (0%, 0.5%, 1%, 3%, 5%, and 10% final concentration (w/v)) in 25 mM Tris-HCl (pH 7.4), 300 mM NaCl, 0.5 mM TCEP. The phase diagram was then plotted according to the images collected from three independent reconstitutions. For reconstitution of LBLs *in vitro*, 10 µM of EGFP-synphilin-1 was mixed with 20 µM α-synuclein (containing 1:10 Alexa Fluor™ 568 C5 Maleimide-labeled α-synuclein) in 25 mM Tris-HCl (pH 7.4), 300 mM NaCl, 0.5 mM TCEP, 10% (w/v) PEG 8000. If not otherwise indicated, the condensation reaction was incubated for 5 min at RT and transferred to a 15-well 3D glass bottom slide (Ibidi) for imaging.

#### Phase separation assay of α-synuclein

For phase separation experiments with α-synuclein, unlabeled protein was mixed with maleimide labeled α-synuclein (1:10 Alexa Fluor™ 568 C5 or Alexa Fluor™ 647 C2) at concentrations of 10 µM or 100 µM, and different percentages of PEG 8000 (0%, 10%, 25%, final concentration (w/v)) in 25 mM Tris-HCl (pH 7.4), 150 mM NaCl, 0.5 mM TCEP. If not otherwise indicated, the condensation reaction was incubated for 5 min at RT and transferred to a 15-well 3D glass bottom slide (Ibidi) for imaging.

### Cell-based phase separation assays

#### Eukaryotic cell lines

Human embryonic kidney (HEK293, ACC 305 from DSMZ; German Collection of Microorganisms and Cell Cultures GmbH) cells were cultured in a humidified incubator at 37 °C with 5% CO2 using Dulbecco’s Modified Eagle Medium (DMEM) containing 4.5 g/L D Glucose (Gibco) supplemented with 10% heat-inactivated FBS (FACS), 1% Penicillin/Streptomycin and 1% MEM medium (non-essential amino acids) (Sigma). When reaching a confluency of 80–90%, cells were passaged using standard trypsinization techniques. For live-cell imaging experiments, HEK293 cells were either grown on glass coverslips (25 mm) or in glass bottom dishes until reaching a confluency of 50–70%.

For recombinant protein purification from mammalian cells the Expi293F^TM^ expression system was used (ThermoFisher Scientific). In brief, Expi293F (#A14527) cells were grown in 30 mL Expi293 expression medium (#A1435101) in suspension (37°C, 8% CO_2,_ 125 rpm) to 5×10^6 cells/mL and subsequently split to ∼0.3×10^6 cells/mL for general maintenance.

#### Modulation of osmotic pressure

HEK293 cells expressing either only EGFP-synphilin-1 or in combination with α-synuclein-BFP were treated with the hypotonic solution (5% DMEM in H_2_O) or hypertonic solution (300 mM NaCl in H_2_O). Images from three independent transfections were collected before and after 5 min from the addition of the solutions.

#### Colocalization of synphilin-1 inclusions with organelles

HEK293 cells expressing either only EGFP-synphilin-1 or in combination with α-synuclein-BFP were incubated with several fluorescent dyes in order to visualize subcellular lipid localization (MM 4-64 (FM 4-64) Dye, ATT Bioquest), lysosomes (LysoTracker™ Deep Red, Thermo Scientific) and mitochondria (MitoTracker™ Deep Red FM, Thermo Fisher). Growth media was replaced by pre-warmed live cell imaging solution (Thermo Fisher Scientific, A14291DJ) containing fluorescent dyes (final concentrations; MM 4-64: 1.65 µM, Lyso and MitoTracker: 1 µM) and cells were incubated for 30 min. Before imaging the staining solution was replaced by fresh live cell imaging solution. The co-localization of synphilin-1 inclusion and LBLs with cellular organelles was analyzed from three independent transfections as a percentage of the overlapping surface area using ImageJ (NIH).

#### Timelapse imaging of synphilin-1 inclusions with cytoskeleton

HEK293 cells expressing EGFP-synphilin-1 and α-synuclein-BFP were incubated with either SPY555-actin (Spirochrome, 1 mM stock solution), SiR-Actin (Spirochrome, 1 mM stock solution), or SiR-Tubulin (Spirochrome, 1 mM stock solution). Fluorescent dyes were diluted to 1 µM (final concentration) in pre-warmed DMEM and incubated with HEK293 cells for 30 min. The staining procedure was initiated at different time points post-transfection (15.5 h or 19.5 h). Before imaging the staining solution was replaced by fresh live cell imaging solution (Thermo Fisher Scientific, A14291DJ) and images were collected every 10 min for 1 hour. For the quantification of actin and tubulin enrichment within the condensates over time a mask of the EGFP-synphilin-1 signal was created. The mean gray intensity value was extracted for the actin or tubulin channel and plotted as a mean ± s.e.m. from three independent biological replicates (at least 5 condensates from each).

#### Latrunculin A treatment in AK-11 cells

AK-11 cells expressing EGFP-synphilin-1 and α-synuclein-Halo were treated with 2.5 µM Latrunculin A (Sigma-Aldrich 428026) 45 min after transfection and induction with 500 ng/mL doxycycline. 23.5 h after transfection, cells were incubated for 30 minutes in 1 mL of their own growth medium supplemented with Janelia Fluor JF549 HaloTag ligand (Promega, HT1020, 1:1000) and actin fluorescent dye (CellMask™ Actin Orange, Thermo Scientific). Before imaging the staining solution was replaced with fresh imaging solution. In addition, untransfected/not treated and transfected/not treated cells were imaged as a control. Images from three independent replicates were collected.

#### Latrunculin A treatment in neuronal cells

Neurons expressing EGFP-synphilin-1 and α-synuclein-BFP were treated with 2.5 µM Latrunculin A (Sigma-Aldrich 428026) immediately after transfection. 24 h after transfection, cells were fixed in 4% PFA (v/v) in PBS, supplemented with 4% sucrose, 1 mM MgCl_2_ and 0.1 mM CaCl_2_, for 15 min at RT and quenched in 50 mM NH_4_Cl in PBS for 30 min, followed by three brief washes with PBS. Cells were then permeabilized for 3 minutes in PBS supplemented with 0.1% of Triton-X (PBS-T). Blocking was performed for 30 minutes in blocking solution, consisting of β-casein solution (Thermo Scientific, 37528) supplemented with 0.1% of Triton-X. Cells were stained with anti-MAP2 (Abcam, ab5392, 1:1000) antibodies diluted in blocking solution for 2 hours at RT. After 3 washes of 10 minutes in PBS-T, cells were stained with secondary antibodies and Phalloidin AF647 (Thermo Scientific, A22287) diluted in blocking solution for 1 hour at RT. Secondary antibodies used were goat anti-rabbit STAR488 (Sigma-Aldrich, 52944, 1:400) and goat anti-guinea pig AF568 (Thermo Scientific, A11075, 1:400). Cells were subsequently washed 3 times for 10 minutes in PBS-T and mounted on glass slides using Fluoroshield medium (Sigma-Aldrich, F6182). Images from at least three independent replicates were collected.

#### Peroxy Orange 1 stainings

HEK293 cells were cultured in 6-well plates in high-glucose DMEM supplemented with 10% FBS and 1% penicillin–streptomycin (P/S). The cells were approximately 60% confluent before transfection. The cells were transfected with hsSNCA-BFP or EGFP-C1-hsSNCAP using jetOPTIMUS® transfection reagent. After 4 h, the transfection medium was replaced with fresh culture medium, and the cells were cultured for an additional 20 h. Subsequently, PO1 was added to the culture medium at a final concentration of 25 µM, and the cells were incubated for 1 h at 37 °C. The cells were then washed three times with prewarmed PBS to remove excess PO1 and imaged using a Leica STELLARIS 8 FALCON confocal microscope (Leica). For PO1 imaging, an excitation wavelength of 543 nm was used, and fluorescence emission was collected from 555 to 575 nm. The acquired images were analyzed using Fiji version 2.14.0, and the resulting data were analyzed using GraphPad Prism 8.

### Quantitative assessment of condensate formation

In this study, we used the intensity of fluorescence from two-dimensional images captured by a confocal microscope as described above, to estimate protein concentrations in droplets. This microscope is designed to take images at a specific focal depth within a layer about 0.9 µm, which approximately matches the diameter of early-stage condensates. We adjusted the fluorescence intensity measurements to compensate for the influence of droplet size, which could otherwise skew our observations of concentration changes over time. For each cell under examination, we captured a sequence of images (known as z stacks) every 15 min across the entire sample. At each time interval, we focused on imaging the central z plane of every monitored condensate, as illustrated in Supplementary Figure 6A.

To calculate the concentration of protein in the condensate: ρ, we computed the total intensity received from the focus layer, by summing the value of all the pixels divided by the area of the condensate on the image.

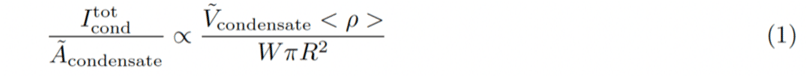

Where I_tot_^Cond^ designate the sum of the intensity value of all the pixels of the 2D image within the droplet region, A_condensate_ and V_condensate_ are respectively the area and the volume of the condensate intersected by the focal plane, < ρ > is the average density of protein in the droplet. Notice that V_condensate_ is a function of the relative position of the droplet with the focus layer, which we do not know. Therefore, we compute the volume intersected, averaged over all the possible position of the droplet relative to the focus layer, provided that the center of the droplet lies within the focus layer. For a given vertical position of the droplet z, we write the volume of interest as:

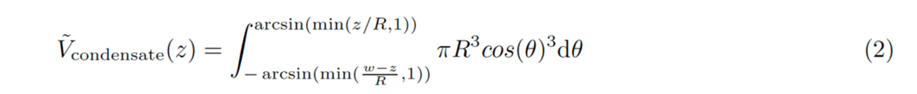

See Supplementary Fig. 6 for notations. Assuming that all positions of the droplet are equiprobable, we write the average volume as:

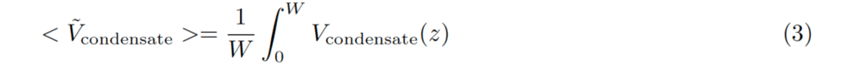

Which gives a different result depending on the relative value of R and W, as illustrated in Supplementary Fig. 6:

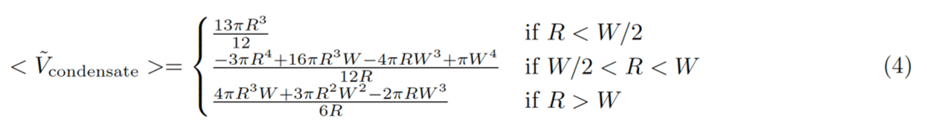

Throughout the study, we assume that transfected cells synthesize synphilin-1 and α-synuclein at a constant rate. To validate this assumption, we plotted here the time evolution of the average total intensity in the cells by summing the value of all the pixels in Supplementary Figure 6a. The linear increase of fluorescence confirms that the production rate is constant throughout the experiment.

To calculate the protein concentration inside the droplet, we applied corrections as described in equations (1)-(4). This correction process starts with accurately determining the radius of the condensates. To determine the radius, we first computed the intensity along a line that crosses the center of mass of the condensate, averaged over all possible angles (also known as radial intensity distribution). We then fit the edge of the average intensity cut using the difference between two sigmoid functions, as depicted in Supplementary Figure 6c, using the fitting function:

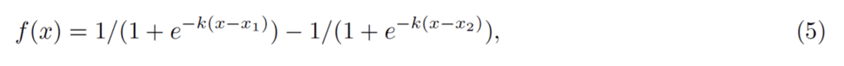

where *k* represents the slope at the boundaries, and x_1_, x_2_ are the positions of the left and right interfaces, respectively. The condensate’s radius is then given as R = (x_2_ − x_1_)/2. Applying this method separately to the synphilin-1 channel and the α-synuclein channel, we obtained the radius changes shown in Supplementary Fig. 6d. We found that α-synuclein is consistently covering an area larger than synphilin-1, even before the visible formation of an α-synuclein shell.

Finally, we plotted in Supplementary Fig. 6e, the corrected intensity ratio in the condensate versus in the cytoplasm for both proteins. An initial decay of fluorescence, that is counterintuitive with the enrichment and growth of the condensate at later stages, can be attributed to inaccuracies in measuring the condensate’s size at the initial stages, due to resolution limit of the confocal microscope. The correction factor for geometrical effects sharply rises with smaller condensates sizes, which amplifies small errors in size estimation. Apart from this very initial stage variation, the patterns shown in Supplementary Fig. 6e guarantee that the initial increase of the averaged intensity in the condensate cannot be only attributed to geometrical artifacts, which fully validate the interpretation of our findings.

### Soft X-ray tomography (SXT)

Prior to SXT imaging, the cells were cryo-fixed through plunge-freezer to preserve the cellular ultra-structures and prevent radiation damage from soft X-ray irradiation. The detailed protocol for preparing cells for SXT imaging was followed^36^. Here, we used rapid plunge-freezing method for the capillary sample holder. Cells were detached from T25 flasks using 0.25% Trypsin and were centrifuged at 300x*g* for 10 min. The cell pellet was mixed with pre-loaded 8 µm microbeads. Next, to prepare the capillary for cryo-fixation, we used a micro-loading tip to load ∼2-3 μL of beads/cells mixture to the pre-cut capillaries with 8-10 μm opening in the tapered tip. The capillary was then rapidly plunged into the liquid nitrogen-cooled liquid propane container for fixation and subsequently stored in the cryogenic storage Dewar until SXT imaging. To transfer the capillaries from storage puck to the microscope, we used the home-built cryo-transfer apparatus purged with liquid-nitrogen-cooled dry helium to ensure the integrity of vitrified cells. Data acquisition for 3D isotropic reconstruction was done by acquiring 2D projections from different angles by rotating the sample stage with every 2-degree increment for 92 projections^89^. Cell segmentation and 3D surface rendering were done by using Fiji (ImageJ) and Amira (Thermo Scientific). The linear absorption coefficient (LAC) and morphological differences were used for identifying different organelles as well as the condensates^37,90^.

#### Quantification of the mitochondrial network

The shape analysis of the mitochondria was done on manually segmented label fields from SXT data. We used the local thickness for the label field as a characteristic thickness and the marching cubes method to determine the surface area^91^. Volumes were defined directly from the sum of the voxels. To account for small defects from noise and/or slice interpolation that the label fields sometimes contain, we applied a Gaussian filter of sigma 1 pixel before the analysis, reducing the amount of artificial surface roughness. Structural characteristics of the mitochondria were determined separately for all disjoint clusters in a cell.

### LC-MS/MS-based proteomic analysis

#### Cell Cultivation and Sample Preparation for LC-MS

Untransfected and uninduced (control) or transfected with EGFP-synphilin-1 and induced for α-synuclein expression AK-11 cells were cultivated as detailed above. 24 h after transfection samples were trypsinized, washed twice in cold PBS and aliquoted into tubes with approximately 100’000 cells per tube. Samples were aspirated, frozen in liquid nitrogen and stored at −80°C.

Proteomics samples were prepared following an adjusted version of the mPOP protocol^92,93^. Briefly, 75 µL of LC-MS grade water was added to samples on ice and samples were incubated at 90°C for 10 minutes in a thermal cycler. Samples were placed on ice immediately afterwards. After they had cooled, 25 µL of lysis buffer were added to each sample, which amounted to a final concentration of 100 mM Triethylammonium bicarbonate (TEAB, Thermo Scientific), 10 ng/µL Trypsin (sequencing grade, Promega), 2 mM magnesium chloride (MGCl2, Roth) and 500 U/10-6 cells Benzonase® nuclease (Merck). The samples were digested for 18 h at 37°C. Digestion was quenched using 20% formic acid (Fisher Chemical) to obtain pH 2-3 and stored at −20°C until measurement.

#### LC–MS/MS Analysis

Based on peptide quantification, 100 ng of peptides were analyzed using an Orbitrap Astral Zoom mass spectrometer coupled to a Vanquish Neo UHPLC system (both Thermo Fisher Scientific). Peptides were separated using a 25 min gradient at a flow rate of 1.25 µL/min with 0.1% formic acid (FA) in water as solvent A and 80% acetonitrile (ACN) containing 0.1% FA as solvent B. The Vanquish Neo system was operated in trap-and-elute mode using a 5.5 cm µPAC column (Neo, 100 – 300 Å; bed dimensions, 1850 µm × 55 mm; pillar dimensions, 2 × 25 µm; maximum pressure, 450 bar; Thermo Fisher Scientific). An EasySpray adapter equipped with a 15 µm inner-diameter emitter was coupled to an EasySpray ion source on the Orbitrap Astral platform (Thermo Fisher Scientific).

Data were acquired in narrow-window data-independent acquisition (nDIA) mode over an m/z range of 380 – 980. Full MS scans were acquired at a resolution of 240,000 (at m/z 200), with an automatic gain control (AGC) target of 500% and a maximum injection time of 3 ms. DIA scans were acquired using a maximum injection time of 3 ms and an AGC target of 75%. Isolation windows with a nominal width of 2 Th were used across the m/z range of 380 – 980. Fragment ions were analyzed in the Astral over a m/z range of 150 – 2000.

#### Protein Identification and Quantification

Raw data were processed using DIA-NN (version 2.3.1 Academia) (Demichev et al., 2019) using a spectral-library-free workflow. Searches were performed against the reviewed human UniProt reference proteome (UP000005640), downloaded on October 10, 2025. DIA-NN was configured with a fragment-ion m/z range of 100 – 1800, mass accuracies of 10 ppm for MS2 and 4 ppm for MS1, a scan window of 6, and Match-Between-Runs (MBR) enabled. Quantification was performed using the “Legacy (direct) quantification mode”. Protein and peptide identification results were filtered to a false discovery rate (FDR) of 1% at the peptide level.

#### Proteomics Data Analysis

Protein abundances were compared between untransfected and uninduced (control) and transfected with EGFP-synphilin-1 and induced for α-synuclein expression (test) AK-11 cells using custom R (for limma differential expression analysis) and Python (for data processing outside of limma analysis and visualization) script routines. Proteins detected in less than 3 out of 5 replicates in both groups were excluded from the further analysis. Differential abundance testing was performed in R using limma package on the non-imputed quantitative protein matrix. log2FC was defined as [mean(log2 abundance in the test group) - mean(log2 abundance in the control group)]. Limma-derived p-values were adjusted for multiple testing using the Benjamini-Hochberg correction, and the resulting adjusted p-values were reported as q-values. For visualization and candidate selection, proteins were considered significantly changed at q ≤ 0.05 and |log2FC| ≥ 0.30 unless otherwise stated. Compartmental annotation was based on Gene Ontology (GO). Precise GO categories used to define compartment membership are listed in Supplementary Table S8. Synphilin-1 and α-synuclein were excluded as intentionally overexpressed proteins from the visualization, but not the analysis, in Figure 4B.

### Electron microscopy

24 h after culturing HEK293 cells on poly-L-lysine coated 6 mm sapphire discs (for cryo fixation) or in culture flasks (correlated light and electron microscopy), cells were transiently transfected with α-synuclein-BFP and EGFP-synphilin-1 using jetPEI® DNA transfection reagent (Polyplus) according to manufacturer’s protocol. Briefly, a ratio of jetPEI®:DNA of 3 µL: 1 µg was used for 24 h. The sapphire discs were covered with a gold-plated 6 mm type-B carrier containing 20% BSA in PHEM buffer, which served as a cryoprotectant. Cells were cryo-fixed using a high-pressure freezer (EM ICE; Leica). For electron tomography of LBLs, freeze substitution was performed in acetone supplemented with 1% osmium tetroxide and 0.5% uranyl acetate and 2% H₂O for 33 h at −90 °C, 14 h at −20 °C and 1h at 20 °C using an automated freeze substitution system (AFS; Leica). Samples were washed with acetone and embedded in epon. Sections (100 nm) were cut using an ultramicrotome (UC6; Leica) and collected on formvar-coated Cu100 grids. Grids were post-stained with uranyl acetate and lead citrate and labeled with 10 nm gold particles. Dual tilt tomograms were recorded by transmission electron microscopy (TALOS 120C; Thermo Scientific) and reconstructed and rendered in 3D using the IMOD software package.

For correlative microscopy, cells were washed with PBS and fixed with 0.2 % glutaraldehyde and 4 % formaldehyde for 20 min at room temperature. Cells were washed twice with PBS containing 0.02 M glycine. Cells were collected by scraping gently in a thin layer of 1 % gelatine. Cells were embedded in 12 % gelatine and small cubes were cryo protected in 2.3 M sucrose in PHEM at 4 °C. 200 nm cryo-sections were cut using an ultramicrotome (UC7/FC7; Leica) and collected on formvar coated and carbon evaporated 100 Mesh nickel grids. Grids were placed in a droplet of 0.1 M PBS pH 7.4 containing 0.33 μg/mL DAPI. Sections were imaged with widefield microscope (Axio Observer Z1; Carl Zeiss) equipped with ZEN software and a Prime 96B digital camera (Photometrics). Z-stack images were acquired using a 100×/1.30 NA objective (Carl Zeiss). GFP fluorescence was detected using a 470/40 nm bandpass excitation filter, a 495 nm dichroic mirror, and a 525/50 nm bandpass emission filter. DAPI fluorescence was detected using a 365 nm excitation filter and a 445/50 nm bandpass emission filter. Sections were washed on bidest water and post stained and embedded in a mixture of 0.5 % uranyl acetate and 0.5 % methylcellulose. Regions of interest identified from fluorescence data were imaged by transmission electron microscopy (TALOS 120C; Thermo Scientific) using MAPS software.

### X-ray nanodiffraction analysis

#### Sample preparation for scanning X-ray nanodiffraction

*Freeze-dried cells* were prepared for scanning X-ray diffraction by chemical fixation followed by plunge-freezing and freeze-drying as previously described^94^. HEK293 cells expressing EGFP-synphilin-1 and α-synuclein-BFP were grown on silicon nitride (Si_3_N_4_) membranes (Silson, Warwickshire, UK) coated with 0.2 mg/mL fibronectin (Sigma-Aldrich, St. Louis, USA) and incubated in PBS for 21 hours before chemical fixation (15 minutes in 4% PFA). Fixed cells were stored at 4°C in PBS. Right before plunge-freezing, the fixed cells were washed with a volatile buffer (85.6 mM triethylammonium acetate in H_2_O) and transferred to the humidity chamber (T=30°C, relative humidity≥84%) of a grid plunger (Leica EM GP, Leica Microsystems, Wetzlar, Germany). The sample was then blotted using standard blotting paper and rapidly plunged into a −195°C cold ethane/propane bath. The frozen-hydrated cells were transferred into a cryo-vial and transported to the vacuum chamber of a custom-built freeze-dryer to remove the remaining amorphous ice by sublimation. The chamber remained sealed for 3 days until the sample reached room temperature. Finally, the freeze-dried cells were transferred into a desiccator and stored in the dark until the X-ray measurements. The samples were monitored by fluorescence and bright-field microscopy at different steps of the preparation procedure (before fixation, after fixation, before vitrification, after freeze-drying). Fluorescence images were acquired using a Zeiss Axio Observer Z1 inverted microscope equipped with an an LD Plan-Neofluar 20x/0.4 objective and Axiocam MRm CCD camera (Zeiss, Jena, Germany), BFP was excited at *λ*=365 nm and EGFP was excited at *λ*=470 nm using suitable bandpass filter sets.

*α-synuclein preformed fibrils (PFFs)* were generated from monomeric alpha-synuclein according to a published protocol^95^ and subsequently stored at −80°C. For the X-ray diffraction measurements, an aliquot of α-synuclein PFFs was thawed, a 10 μL droplet was pipetted onto a silicon nitride membrane (Silson, Warwickshire, UK), and air-dried. The sample was then transferred into a desiccator and stored in vacuum until the X-ray measurements.

*Human brain tissue* from an individual who underwent diagnostic autopsy in the context of routine clinical care was obtained from the archives of the Department of Neuropathology UMG in accordance with UMG ethics regulations. Small brain tissue blocks were dissected from 10% formalin-fixed brain slices, dehydrated, and paraffin embedded^96^. 10 μm thick sections of the formalin-fixed paraffin-embedded (FFPE) tissue block were cut using a microtome and deposited on 1.4 μm thin PET membranes (MMI, Eching, Germany). For optical microscopy, the sections were deparaffinized, rehydrated, and H&E stained. Whole-slide images were acquired using a VS200 microscope equipped with a 10x or 20x UPLXAPO objective and operated by the cellSense Dimension software (Evident Corporation, Tokyo, Japan). Images were stored and annotated using an OMERO server^97^. Lewy bodies in a section of the substantia nigra were identified by a trained neuropathologist (J.F.). The sample was stored in water at 4°C until the X-ray measurements. For X-ray nanodiffraction analysis at the ESRF, the section was air-dried and the metal frame of the slide was glued to a brass pin for mounting. After the X-ray measurements, further optical micrographs of the dried section were taken using the VS200 microscope.

#### Scanning X-ray nanodiffraction experiments

Experiments were carried out at the nanofocus endstation (EH3) of beamline ID13 at the European Synchrotron Radiation Facility (ESRF). The beamline configurations are summarized in Supplementary Table S4. An X-ray beam of 15 keV photon energy was focused by multilayer Laue lenses to a 250×250 nm^2^ spot at the sample plane (FWHM). The sample was mounted onto a sample stage consisting of a hexapod and a high-precision piezoelectric stage for accurate lateral scanning. An on-axis optical microscope was used for positioning the sample and selecting the scan regions after comparison to the optical micrographs taken prior to nanodiffraction experiment. The sample was then raster scanned (250 nm horizontal step size and 500 nm vertical step size between subsequent lines) through the focus of the X-ray beam, illuminated for 5 ms at each scan point, and the far-field scattering pattern was recorded by an Eiger 4M hybrid photon-counting detector with a pixel size of 75 μm (Dectris, Baden, Switzerland) located ∼0.34 m behind the sample. The primary beam was blocked by a lead beamstop. The scanning speed and the exposure time were chosen based on test measurements (data not shown), such that radiation damage is not yet visible in the diffraction signal. All samples were measured in a dry state to prevent secondary damage due to water radiolysis and the spread of highly reactive byproducts.

#### Analysis of the scanning X-ray nanodiffraction data

The two-dimensional (2D) detector images were normalized by the primary beam intensity, as measured by an ionization chamber located upstream of the sample, to account for changes in the photon flux during a scan. The images were then azimuthally averaged using a custom Python program based on pyFAI^98^ and the silx toolkit^99^, yielding a one-dimensional (1D) scattering profile at each scan point. Detector gaps, bad pixels and the shadow of the beamstop on the detector were masked before azimuthal averaging, and the data between *q*=[1.81, 1.835] Å^-1^ were excluded from further analysis due to a narrow peak at q=1.82 Å^-1^ that was present in the instrumental background, presumably due to residual scattering from a platinum aperture before the sample. X-ray dark-field maps were calculated by integrating the intensity (number of scattered photons) in the *q*-range between 0.4 Å^−1^ < *q* < 1.9 Å^−1^, where *q* is the magnitude of the scattering vector defined as 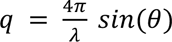, 2*θ* is the scattering angle and *λ* is the wavelength. The regions of interest (ROIs) within the sample and the background regions (containing little to no sample material) were selected semi-correlatively by comparing the X-ray dark-field maps to the optical micrographs of the sample. The 1D scattering profiles were averaged over the scan points within a ROI yielding the ROI average *I_R_*_0*I*_(*q*). The mean scattering intensity of a background region *I_back_*_g*round*_(*q*) was subtracted from *I_R_*_0*I*_(*q*) to get the background-corrected scattering intensity profile of the sample within the ROI. To determine the peak positions and intensities, a model function *(1)* consisting of the sum of three Gaussians plus a constant baseline was fitted to the background-corrected scattering profile in the range between *q* = [0.4, 1.9] Å^-1^ using the Levenberg-Marquardt algorithm.

Fit function: sum of three Gaussians plus a constant:

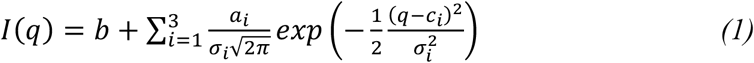

where *b* is a constant baseline, *c_i_* is the center of the Gaussian peak, *a_i_* is the area under the Gaussian peak, and *σ_i_* is the width parameter which is related to the full width at half maximum (*FWHM*):

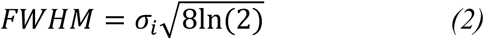

The height of the Gaussian peak *h_i_* can be calculated as:

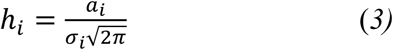

## QUANTIFICATION AND STATISTICAL ANALYSIS

The sample sizes for all experiments were determined by the culture, observation field and ROI. No data were excluded from analysis. All attempts at replication were successful. The study did not use group allocation. For the *in vitro* part, blinding was not typically used. In fact, the proteins had to be separately purified with different protocols and the buffer conditions of different groups need to be carefully controlled by the experimentalists. For the key cell imaging experiments, we have done double-blind transfection by one experimentalist and the imaging by another experimentalist, to ensure the reproducibility. Details on statistical analysis for each individual experiment could be found in figure legends and in method details section.

## ACKNOWLEDGMENTS

We thank for support the Advanced Medical Bioimaging Core Facility and High Throughput Mass Spectrometry Core Facility (funded under the Excellence Strategy of the Federal Government and the Länder by the Berlin University Alliance) at Charité University Clinic, Berlin. The work is supported by the ERC Grant MemLessInterface (101078172), the Horizons Europe MSCA DN Grant (101167843 - BICEPS) to DM and GB; the grants from the German Research Foundation (MI 2104 and SFB1286 Quantitative Synaptology Projects B10 to DM; A02 to TS; A03/B02 to SOR; B08 to TFO; Innovative Minds Program to CH; Alzheimer’s Association (AARG-25-1486936) to YD. JVT is supported by a stipend from the Sonnenfeld Foundation. The US NIH R01NS121114 to RVP and the Mark Foundation for Cancer Research ASPIRE grant to RVP; and Center for Biomolecular Condensates in the James McKelvey School of Engineering at Washington University to RVP and KMR. The National Center for X-ray Tomography is supported by NIH NIGMS (P41GM103445 and P30GM138441) and the DOE’s Office of Biological and Environmental Research (DE-AC02-5CH11231). The Center is located at the Advanced Light Source, a U.S. DOE Office of Science User Facility.

## AUTHOR CONTRIBUTIONS

Conceptualization: DM

Investigation and Visualization: RS and CH did all reconstitution and cell lines experiments and analyzed the data. AK, PB, GS performed longitudinal live-cell imaging, immunostainings in cell lines and data analysis. JHC, AE, MLG and CL performed soft X-ray tomography and analyzed the data. AK, KL, KTT, VD, MM and MR performed proteomics dataset acquisition. AK established the AK-11 cell line. AK, GAP, HW and AN performed experiments in neurons and analyzed the data. AJE and BJE contributed to experimental planning and discussion. AN and PG performed analysis of human samples. RB, LTH and GB performed CLEM experiments and analysis. JJ and DB performed the assessment of mitochondria fitness. MC and YD did PO1 experiments and data analysis. AM and TS performed X-ray nanodiffraction experiments and analysis. JF and CS contributed to X-ray nanodiffraction experiments in tissues.

Formal Analysis: HLR and PDLR did quantitative assessment of phase separation and modeling. KR and RP did bioinformatics modeling of molecular grammars. KL, KTT, AK, VD, LS, MM and MR performed the analysis of proteomics datasets.

Resources support: SOR, TFO, DS, MR, DS, BJE

Writing – original draft: RS, CH, AK, JHC, CL and DM.

Writing – review and editing: all contributing authors.

Funding acquisition: DM Project administration: DM

---

**Supplementary Figure 1.**
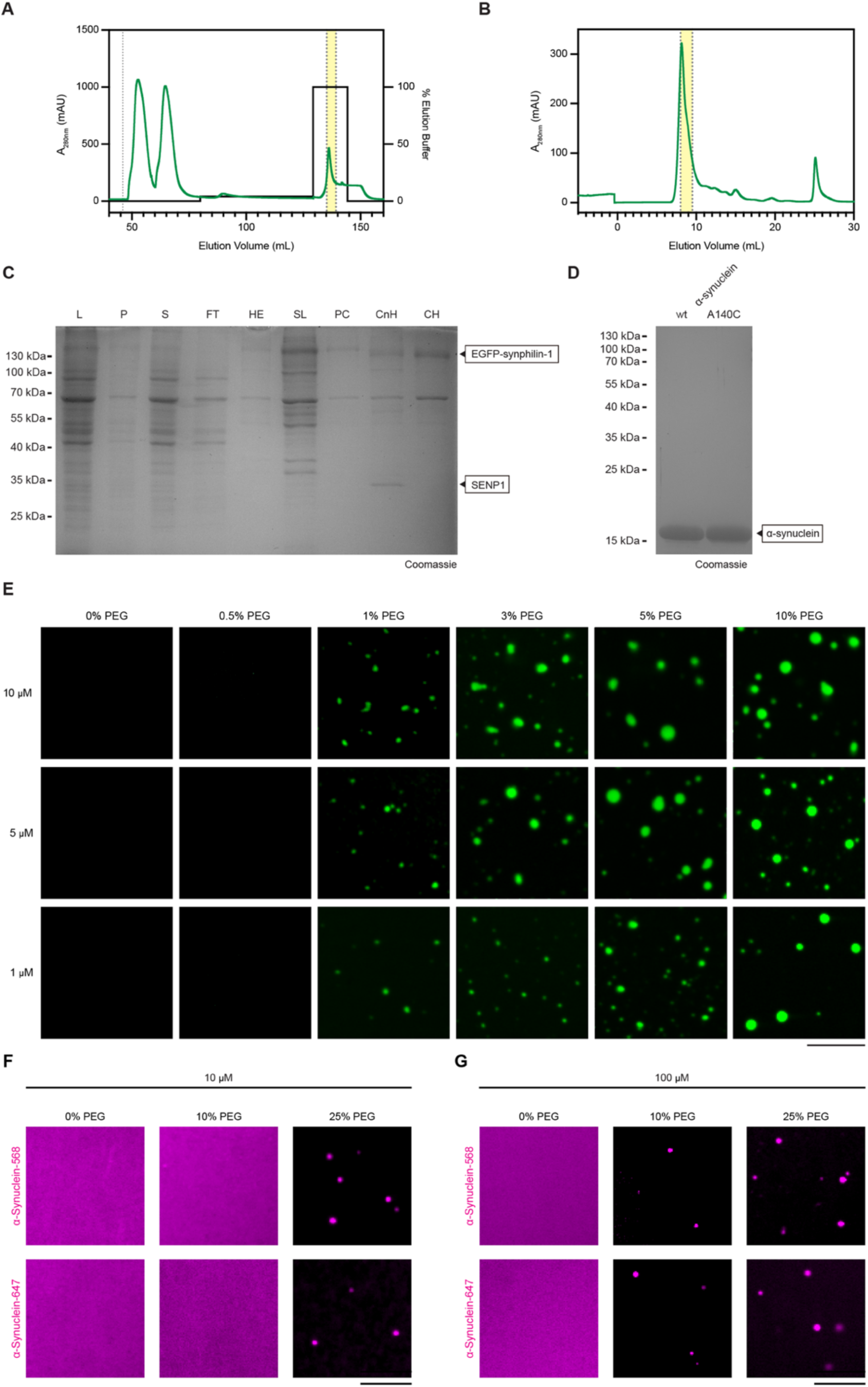
Recombinantly purified synphilin-1 undergoes phase separation while α-synuclein remains soluble under physiological salt concentration and crowding. **(A)** Chromatogram showing affinity purification of His-SUMO-EGFP-synphilin-1. Absorbance at 280 nm (y-axis, left) and the buffer employed expressed as percentage of elution buffer (y-axis, right). The fractions collected are highlighted in yellow. **(B)** Chromatogram showing size-exclusion (S200) purification of His-SUMO-EGFP-synphilin-1. The fractions collected are highlighted in yellow. **(C)** SDS-page gel showing all steps of EGFP-synphilin-1 purification (L: cell lysate, P: pellet of cleared lysate, S: supernatant of cleared lysate, FT: flow-through of affinity column, HE: His-eluate, SL: load to size exclusion column, PC: Pre-cleavage, CnH: SENP1 cleavage reaction, CH: reverse Ni-NTA flow-through). **(D)** SDS-page gel showing the recombinant, purified proteins used in the reconstitution assays: α-synuclein wild-type (left) and α-synuclein A140C (right). **(E)** Representative images of EGFP-synphilin-1 phase separation at different concentrations: 1 μM, 5 μM, 10 μM, and with increasing percentage of PEG 8,000 (0%, 0.5%, 1%, 3%, 5%, 10% w/v). Reconstitution experiments were performed at physiological salt concentrations: 25 mM Tris-HCl (pH 7.4), 300 mM NaCl, 0.5 mM TCEP. Samples were imaged at ambient temperature 5 min after incubation. **(F)** Reconstitution of 10 μM α-synuclein (chemically labeled with Alexa Fluor 568 or Alexa Fluor 647 (α-synuclein-568 and α-synuclein-647)) in varying percentages of PEG 8,000 (0%, 10%, 25% w/v). **(G)** Same as in *F* but using 100 μM α-synuclein. Scale bars, 10 μm. All data were consistent in three independent reconstitutions.

**Supplementary Figure 2.**
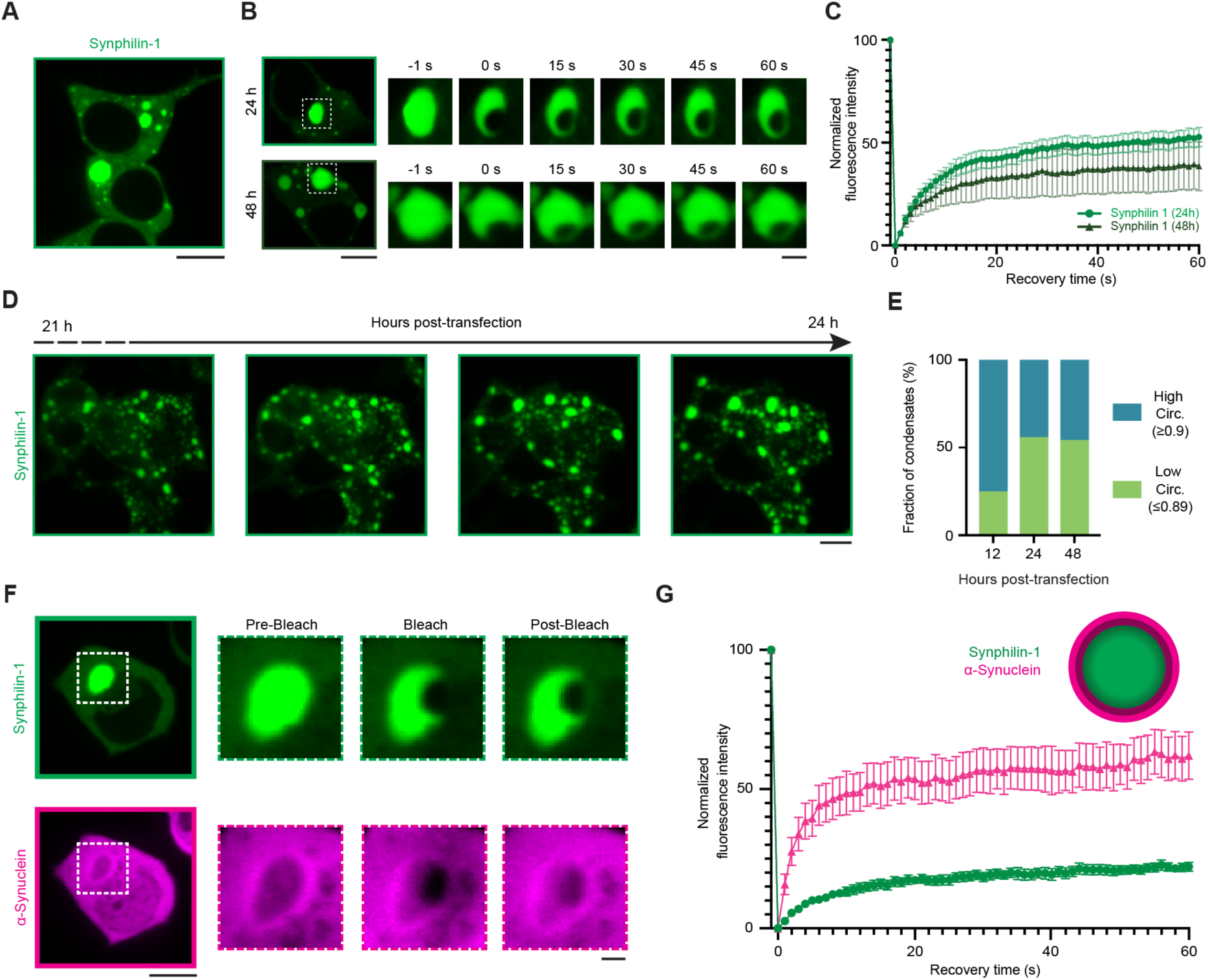
Synphilin-1 undergoes crowder-mediated phase separation. **(A)** Confocal image of EGFP-synphilin-1 ectopically expressed in HEK293 cells. Scale bar, 10 μm. **(B)** Representative fluorescence images of EGFP-synphilin-1 at 24 h (top) and 48 h (bottom) post-transfection in HEK293 cells before, during bleaching, and upon recovery after 60 s. Scale bars, 10 μm (overview), 3 μm (magnified). **(C)** Fluorescence recovery after bleaching of EGFP-synphilin-1 24- and 48-hours post-transfection in HEK293 cells. Values represent average ± standard error of the mean. Data are from three independent replicates. **(D)** Time-course images of cells expressing EGFP-synphilin-1, showing multiple fusions events leading to the formation of larger condensates. Scale bars, 10 μm. **(E)** Histogram showing the fraction of EGFP-synphilin-1 condensates exhibiting low circularity (≤0.89) and high circularity (≥0.9) at three time points (12 h, 24 h, and 48 h) post-transfection. Data from three independent replicates. **(F)** Co-expression of EGFP-synphilin-1 and α-synuclein-BFP results in demixing and accumulation of α-synuclein in the shell and synphilin-1 in the core. Right: magnification of EGFP-synphilin-1 and α-synuclein-BFP before and during bleaching, and upon recovery after 60 s (post-bleach). Scale bars, 10 μm (overview), 2 μm (magnified). **(G)** Fluorescence recovery after photobleaching of EGFP-synphilin-1 (green) and α-synuclein-BFP (magenta) at the rim of condensates. Values are represented as average ± standard error of the mean. The experiment was performed in three independent replicates, with at least ten condensates analyzed.

**Supplementary Figure 3.**
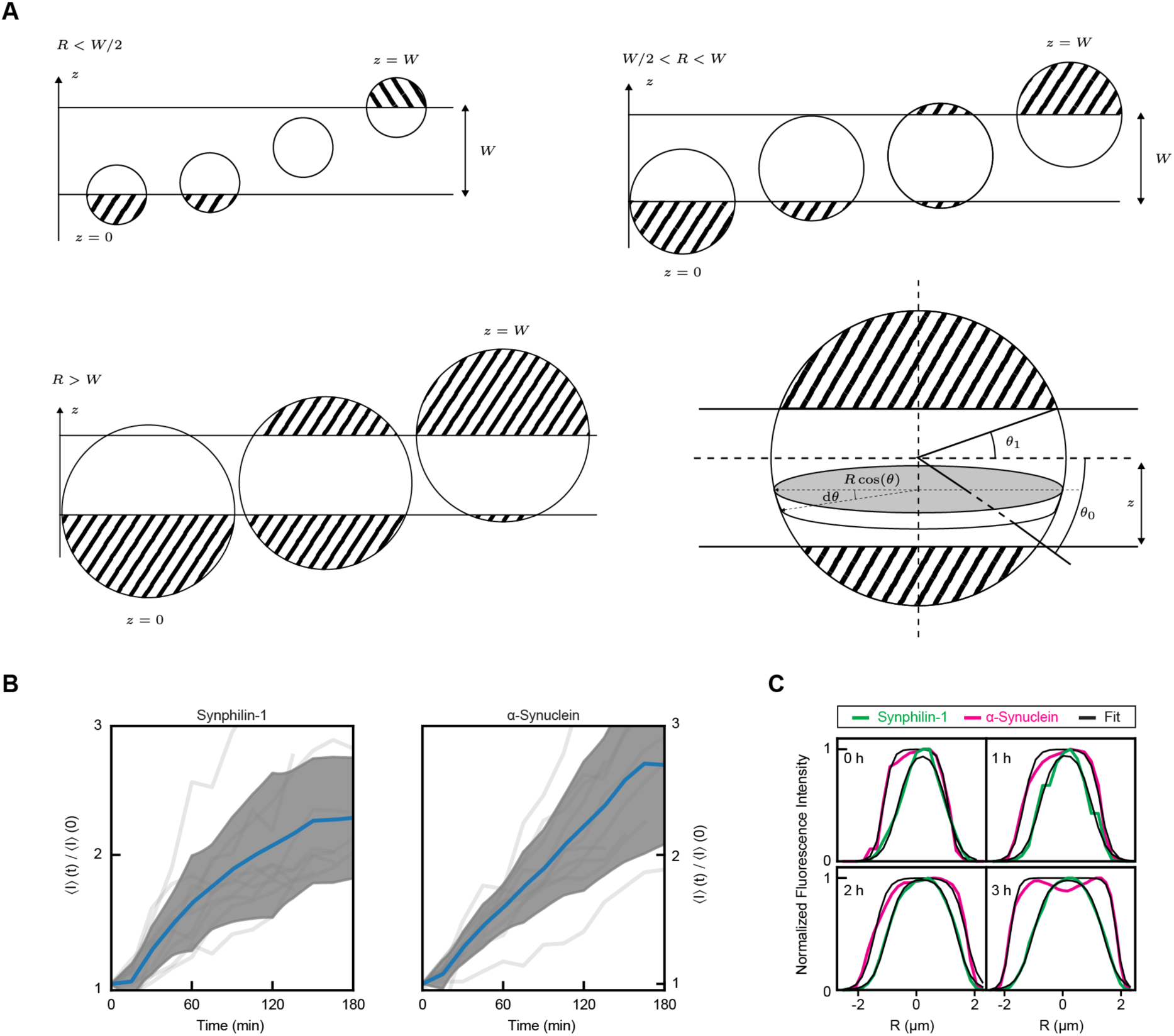
Computation and modeling of the fluorescence intensity evolution in cells expressing Lewy body-like condensates. **(A)** Schematic of the volume intersected by the focus layer as a function of the position of the droplet for different regimes of the droplet radius. The dashed part of the droplet corresponds to the portion of the condensate that is not included in the focus layer. We assume that the droplet can be at any depth relative to the focus layer, from z = 0 to z = W, equiprobably, to compute the average volume. The volume is computed by summing infinitesimal slices of the sphere from θ_0_ to θ_1_. **(B)** Change in the relative concentration of proteins (synphilin-1, left; α-synuclein, right) over time. Individual cells are represented by the gray curves, while the blue line denotes the average. The shaded gray area indicates a range of two standard deviations. **(C)** Representative fits of the radial intensity profiles of the condensate boundary using the difference of two sigmoid functions to extract droplet radius measurements at selected time points.

**Supplementary Figure 4.**
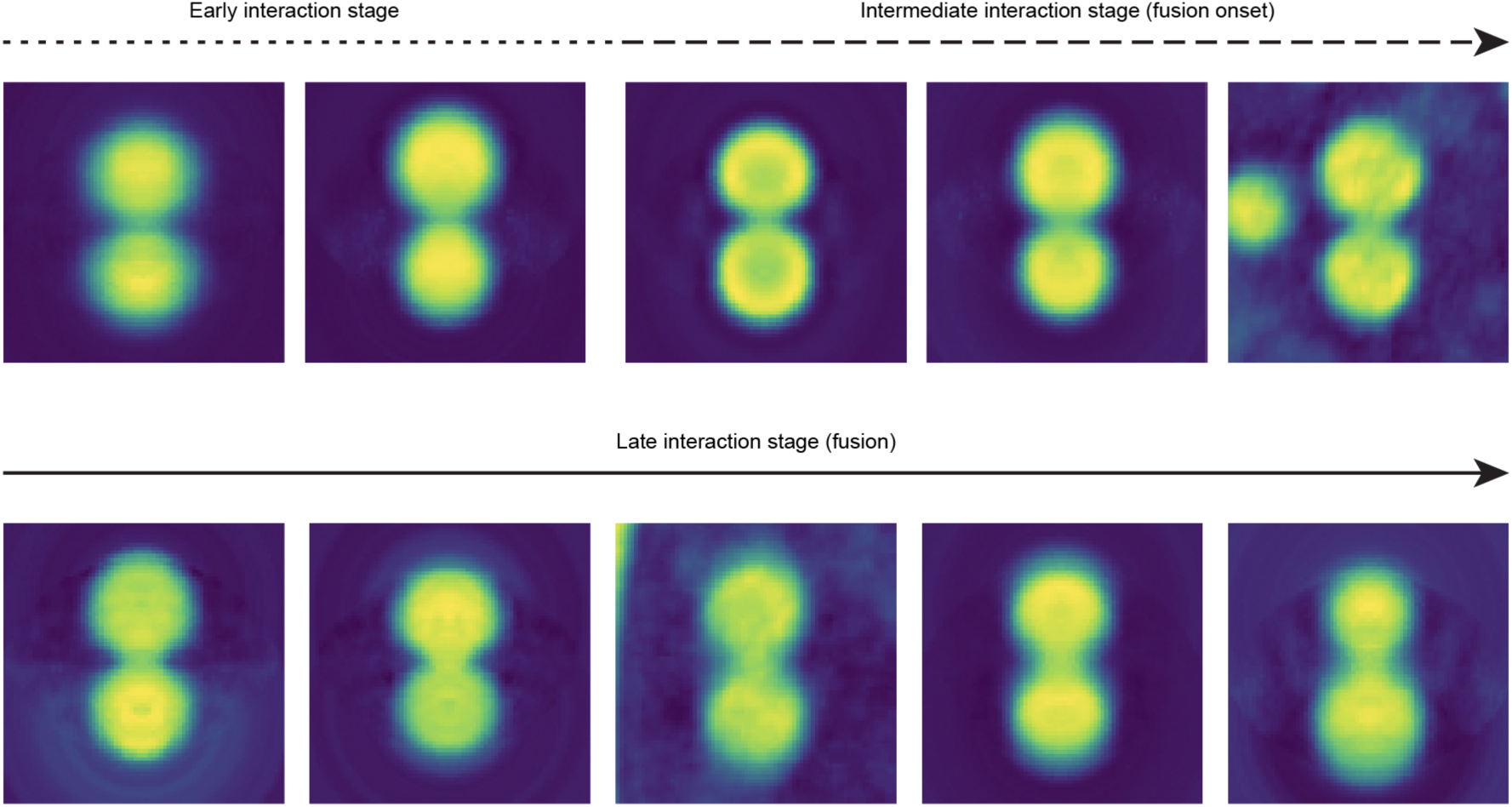
Soft X-ray tomography of Lewy body-like condensates fusion process. Scale-free interpolation of binned histograms based on the linear absorption coefficients (LAC) from LBLs indicates early, intermediate, and the late interaction stages. The cross sections were from the middle slice of the reconstructed virtual slice. Each image is a separate pair of condensates.

**Supplementary Figure 5.**
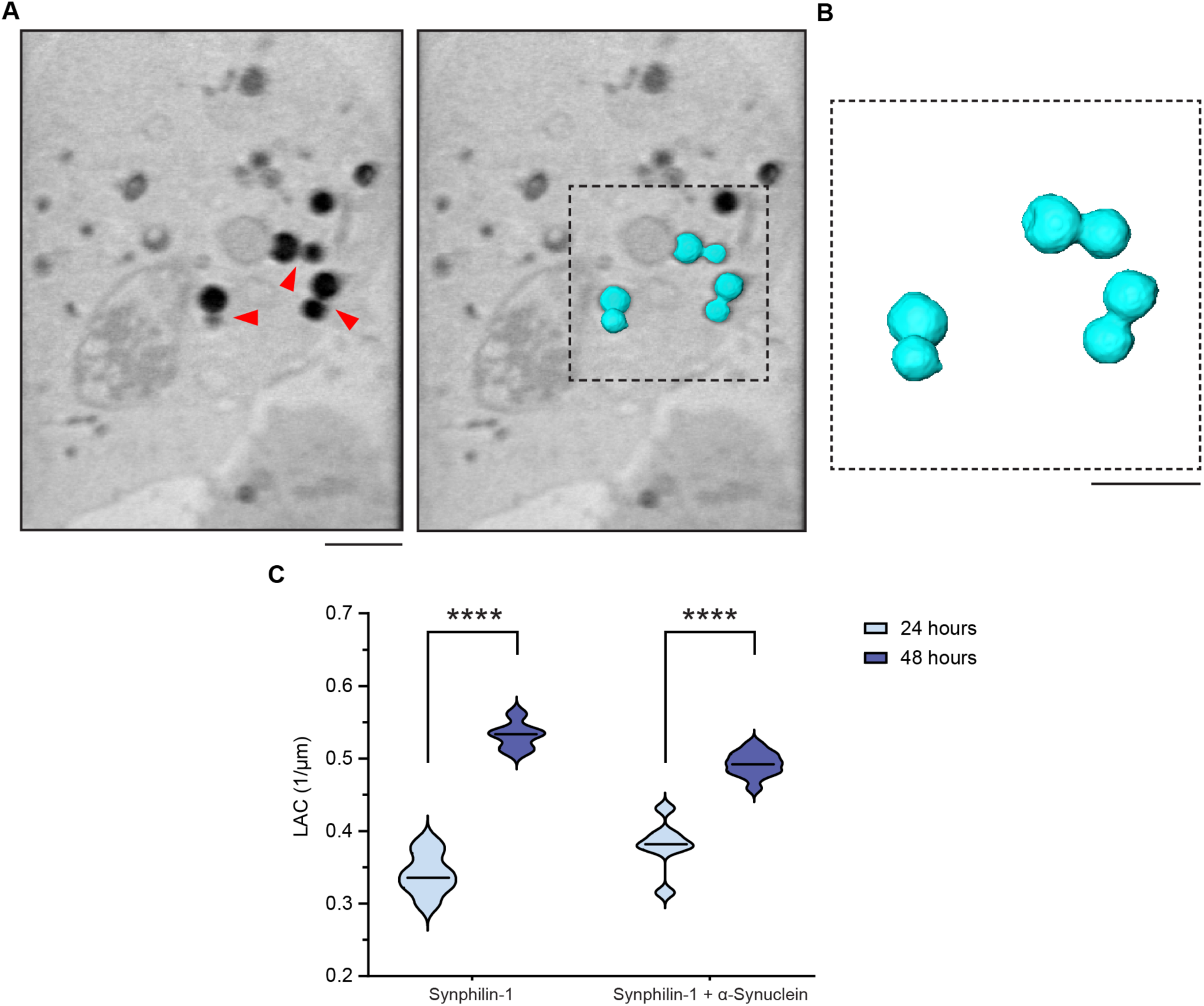
Soft X-ray tomography (SXT) demonstrates the maturation of Lewy body-like condensates. **(A)** Left: orthoslice from a representative tomogram of a cell expressing LBLs (EGFP-synphilin-1 and α-synuclein-BFP). Red arrows indicate LBLs. Right: highlighted surface-rendered LBLs (cyan). Scale bar, 2 μm. **(B)** Surface-rendered LBLs of a selected region in *A* (red arrowheads) captures fusing LBLs. Scale bar, 2 μm. **(C)** Quantification of linear absorption coefficients (LACs) for synphilin-1 condensates and LBLs suggests a change in the material properties (i.e., an increase in density of carbon-rich molecules) in condensates over time; three independent experiments. p ≤ 0.0001, unpaired *t*-test.

**Supplementary Figure 6.**
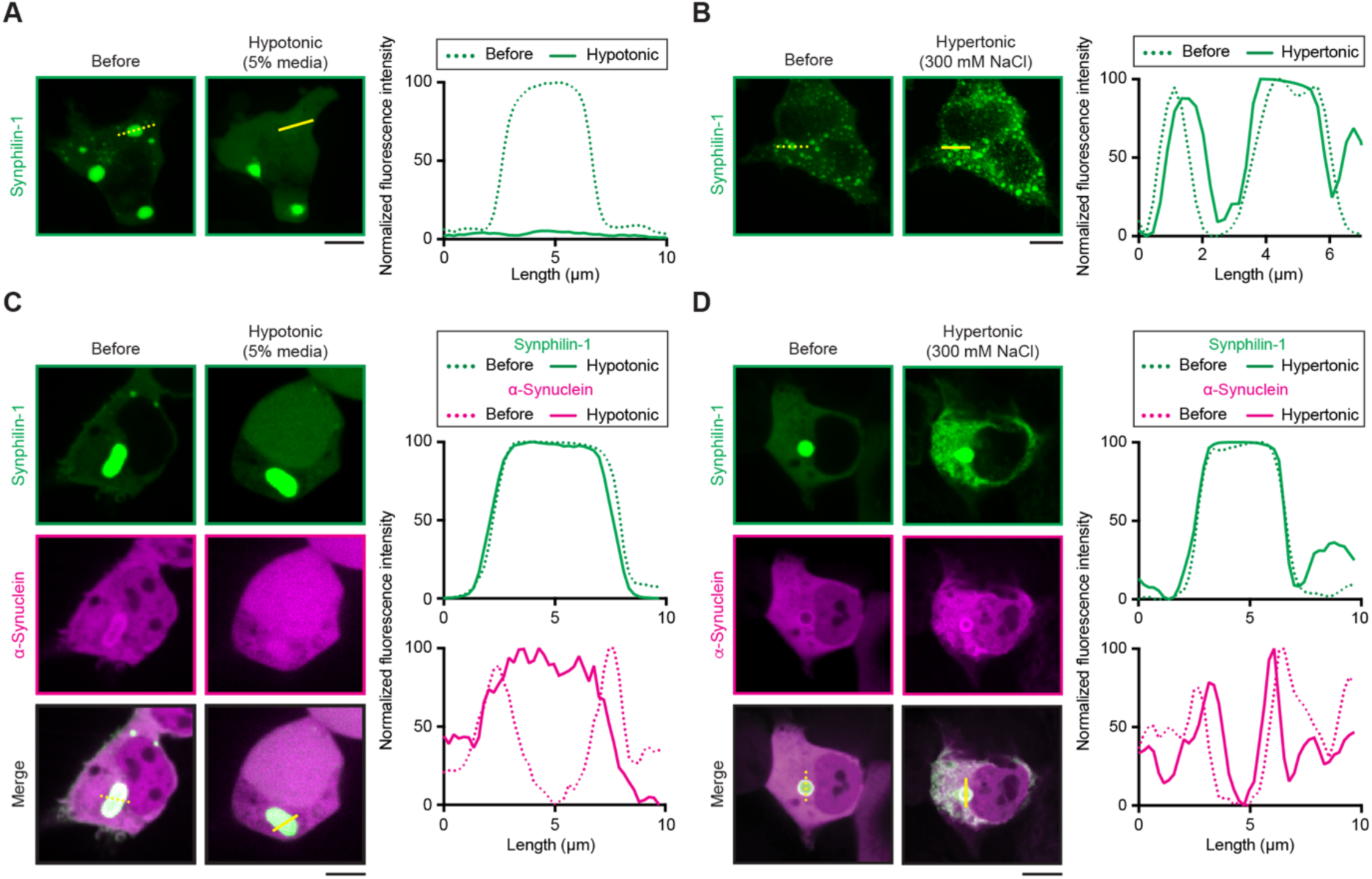
The hypotonic treatment leads to mixing of α-synuclein shell with the EGFP-synphilin-1 core. **(A)** Left: confocal images of HEK293 cells expressing EGFP-synphilin-1, before and after the addition of 5% DMEM in H_2_O. Right: Fluorescence intensity profiles before (solid line) and after (dotted line) exchange of media. **(B)** Same as in *A* but before and upon perfusion of 300 mM NaCl solution (hypertonic solution). **(C)** Left: confocal images of cells expressing EGFP-synphilin-1 and α-synuclein-BFP, before and after exchange of live cell imaging media to the hypotonic solution (5% DMEM in H_2_O); Right: Line profiles for synphilin-1 (green) and α-synuclein (magenta). Fluorescence intensity profiles in conditions before (solid lines) and after (dotted lines) treatment. **(D)** Same as in *C* but before and after perfusion with hypertonic solution (300 mM NaCl in H_2_O). Scale bars, 10 μm. All experimental conditions were probed in three independent replicates. Yellow lines indicate lines along which fluorescence intensity profiles were obtained.

**Supplementary Figure 7.**
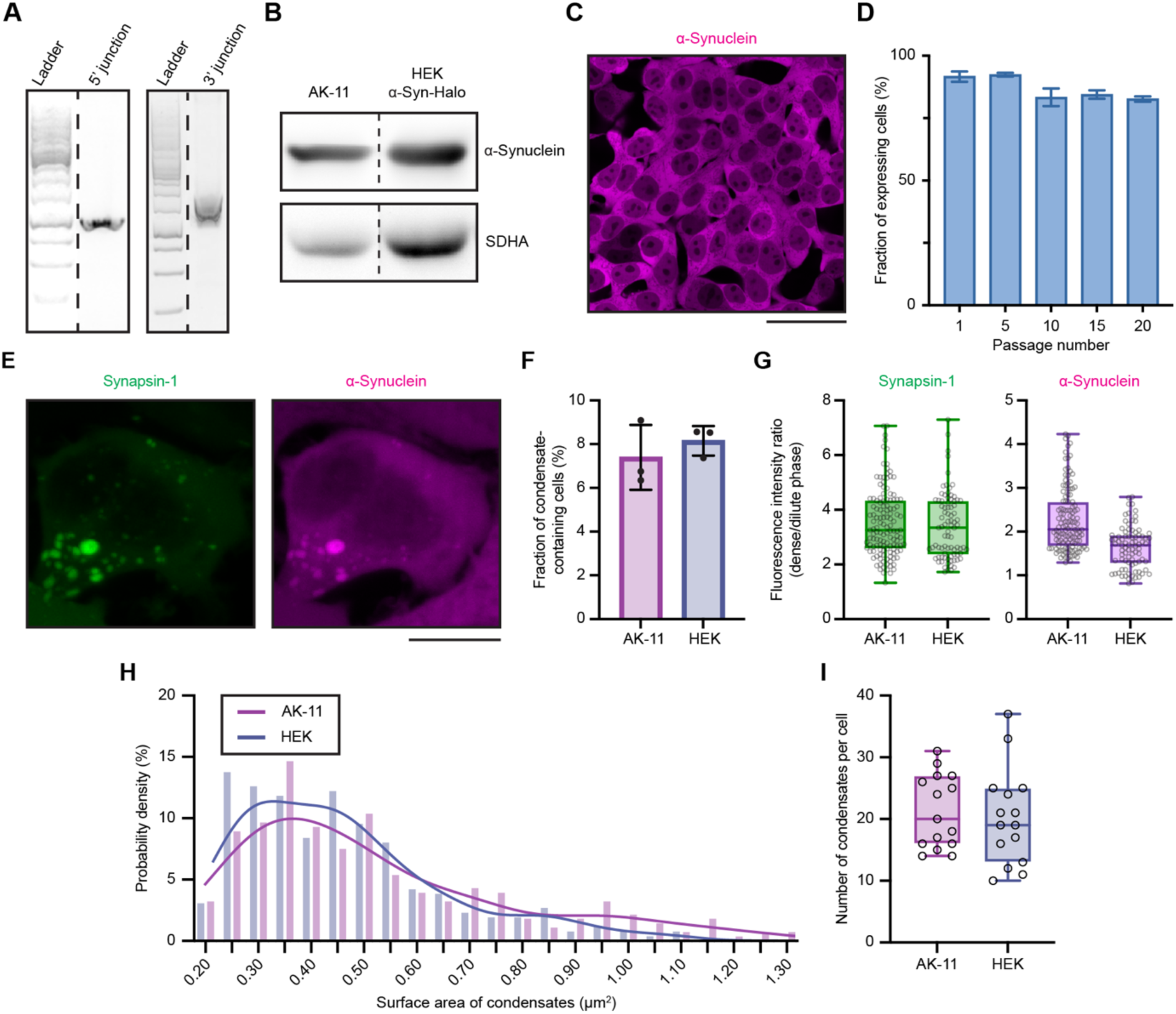
Validation and characterization of the α-synuclein-Halo inducible expressing cell line (AK-11). **(A)** Agarose gel electrophoresis for PCR products of junctions between the integration cassette and the genomic part of AAVS1 integration locus in the AK-11 cell line. **(B)** Western blots of α-synuclein expression level in the AK-11 cell line and after transient transfection in HEK293 cells. Lysates were collected 24 hours after induction or transfection respectively and immunoblotted against α-synuclein. Dashed lines indicate spliced irrelevant bands. **(C)** Representative confocal image of AK-11 cell line in culture. Scale bar, 50 µm. **(D)** Changes in the fraction of expressing cells over passages. AK-11 cells are stable throughout time. Data from at least 1300 cells across three independent replicates for each timepoint. **(E)** Representative images of AK-11 cells, transfected with mCherry-synapsin-1 in AK-11 simultaneously with the induction of α-synuclein expression. Scale bar, 10 µm. **(F)** Fraction of transfected cells that contain condensates after mCherry-synapsin-1 transfection in AK-11 and mCherry-synapsin-1 and α-synuclein-BFP co-transfection in HEK293 cells. Data from at least 600 cells across three independent replicates. **(F)** Synapsin-1 (left) and α-synuclein (right) enrichment in condensates imaged 24 hours after mCherry-synapsin-1 transfection and induction in AK-11 and after double transfection in HEK293 cells. **(H)** Synapsin-1/α-synuclein condensates size frequency distribution 24 hours after mCherry-synapsin-1 transfection and induction in AK-11 and after double transfection in HEK293 cells. Lines represent kernel distribution estimate curves. **(I)** Number of condensates per cell 24 hours after mCherry-synapsin-1 transfection and induction in AK-11 and after double transfection in HEK293 cells. In *G* and *I* the box stretches from the 25th to 75th percentile, dots represent individual data points, the central line shows the median, and whiskers represent the min and max values. Data in *G*, *H* and *I* were collected from at least 15 cells across three independent replicates.

**Supplementary Figure 8.**
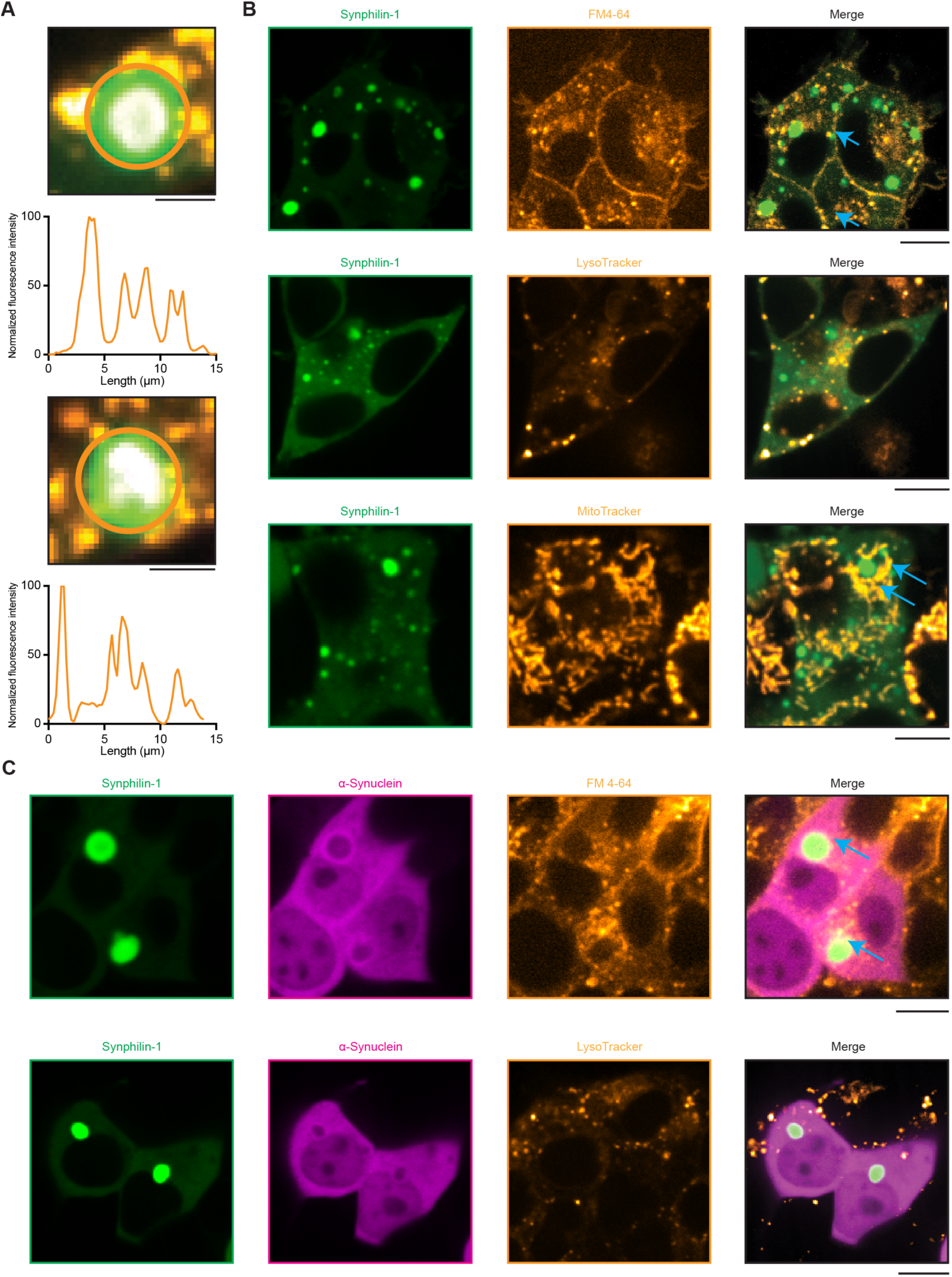
Membrane-bound organelles accumulate in the vicinity of Lewy body-like condensates. **(A)** Images of two representative condensates, each from a separate experiment, of cells expressing LBLs (green, EGFP-synphilin-1) and incubated with MitoTracker-647 (orange) and SPY555-actin (white in overlay) and corresponding circular line profiles. Orange lines indicate the line along which circular profile was obtained in the respective image. Scale bars, 3 µm. **(B)** Representative images of cells expressing EGFP-synphilin-1 incubated with FM4-64 (top), LysoTracker-647 (middle) and MitoTracker-647 (bottom) for 30 min before imaging. Blue arrows indicate organelles in the vicinity of LBLs. Scale bars, 10 μm. **(C)** Representative images of cells expressing EGFP-synphilin-1 and α- synuclein-BFP incubated with FM4-64 (top) and LysoTracker-647 (bottom) for 30 min before imaging. Blue arrows indicate organelles in the vicinity of LBLs. Scale bars, 10 μm. All experimental conditions were performed in three independent replicates.

**Supplementary Figure 9.**
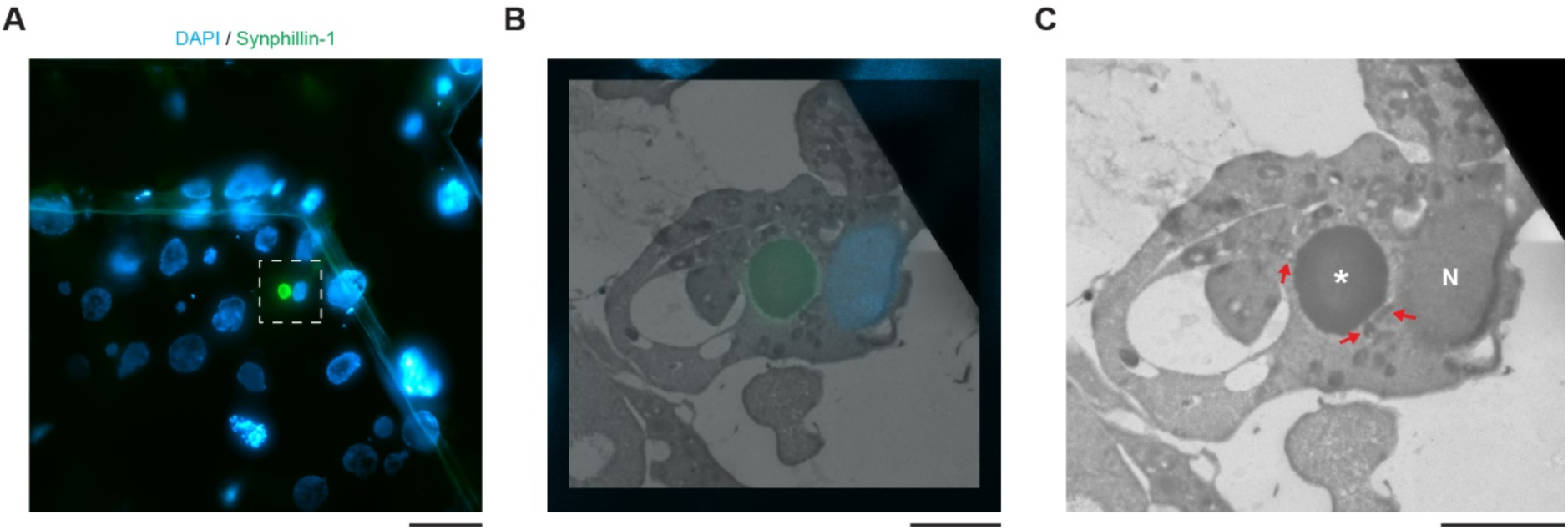
Correlative light-electron microscopy of Lewy body-like condensates expressed in HEK cells. **(A)** Representative overview of HEK cells, expressing EGFP-synphilin-1 and α -synuclein-BFP fixed on EM grid. White dashed line indicates outline of zoomed region. Scale bar, 20 μm. **(B)** Magnified CLEM image of individual cell from *A*. Scale bar, 5 μm. **(C)** Transmission electron microscopy image for the cell from *B*. Red arrows indicate organelles in close proximity of LBL. N, nucleus; *, LBL. Scale bar, 5 μm.

**Supplementary Figure 10.**
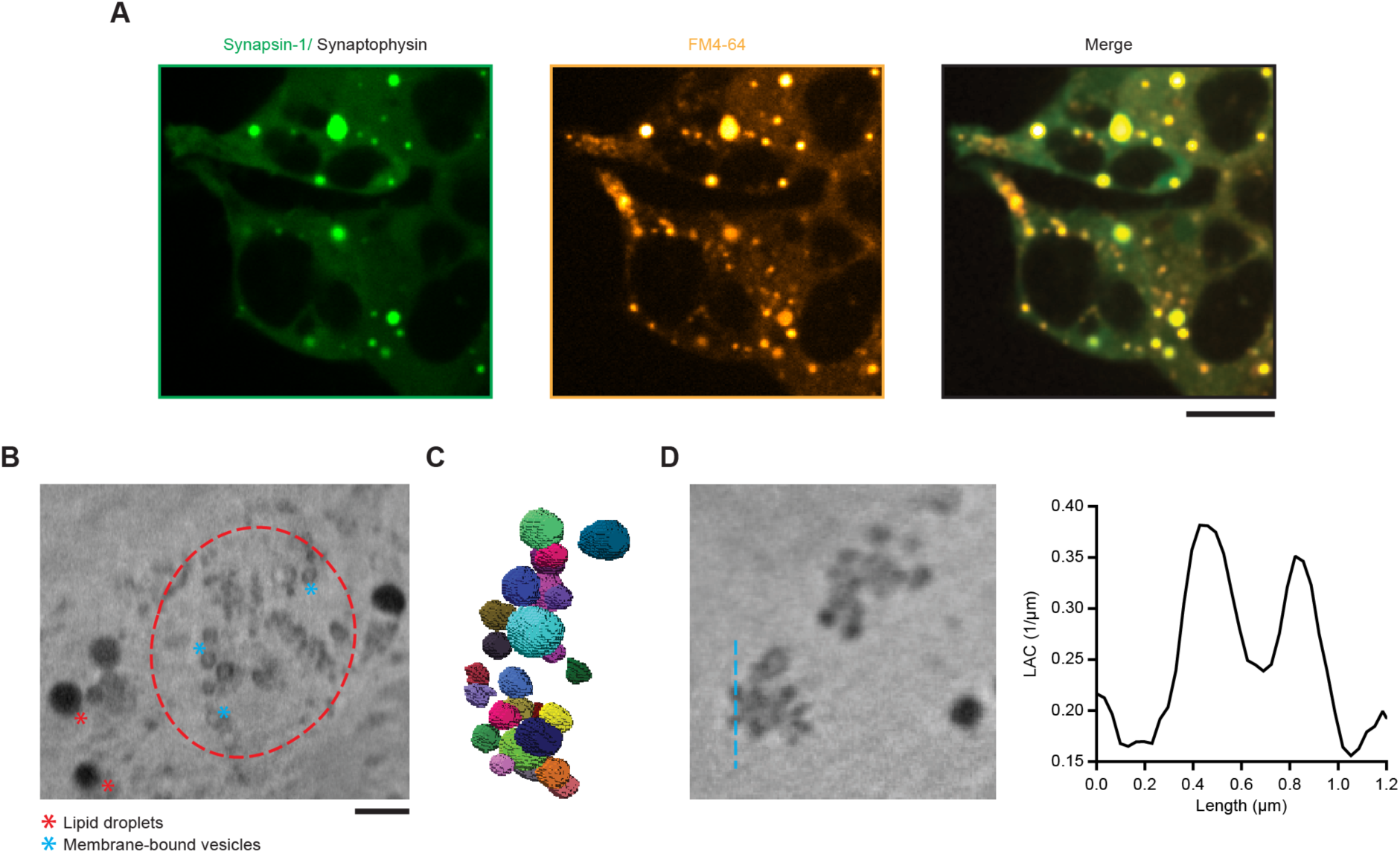
Characterization of ectopically-formed synaptic vesicle-like condensates by soft X-ray tomography. **(A)** Representative images of HEK293 cells expressing EGFP-synapsin-1 and untagged synaptophysin, incubated with FM4-64 (membrane marker). Note the formation of large circular condensates enriched with membranes. The experiment was performed in three independent replicates. Scale bars, 10 μm. **(B)** Orthoslice (virtual section) of a representative tomogram of a cell expressing EGFP-synapsin-1 and untagged synaptophysin. Red dashed line indicates cohorts of small vesicles. Scale bar, 3 μm. **(C**) Surface-rendered view of a condensate of membrane-bound vesicles inside the cell in *B*. **(D)** Left: A representative orthoslice of a tomogram through vesicular cohorts. Right: Line profile through the equatorial plane of a vesicular condensate, indicating the linear absorption coefficient (LAC). Note that each vesicular structure can be clearly separated based on LAC values.

**Supplementary Figure 11.**
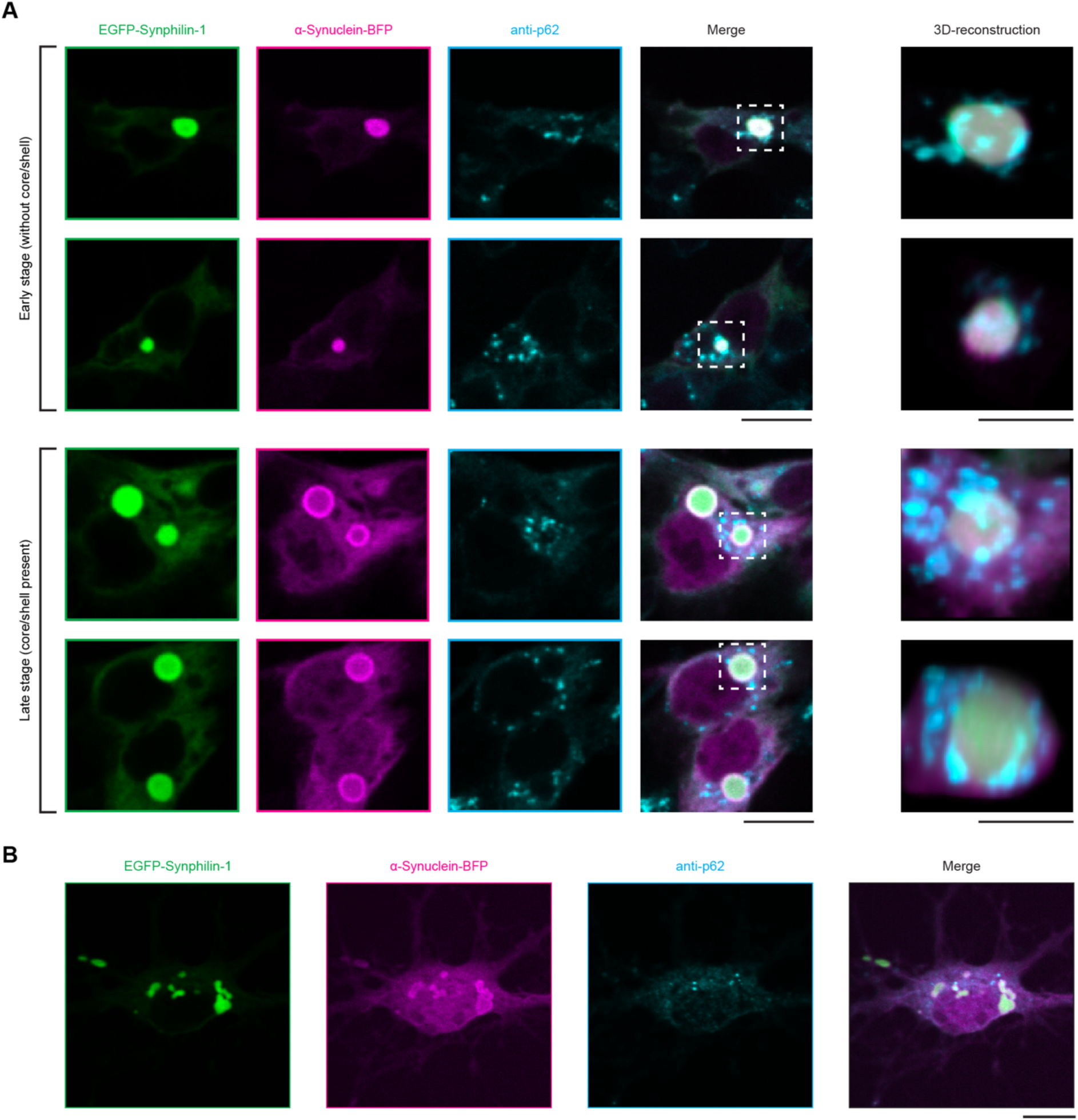
Lewy body-like condensates recruit p62 bodies, consistent with autophagic cargo recognition. **(A)** Immunostaining for p62 in HEK293 cells expressing EGFP-synphilin-1 and α-synuclein-BFP in early stages with fully mixed signal of LBLs (top) and the mature LBLs with core-shell architecture (bottom). Note the enrichment of p62 puncta around LBLs in both cases. Scale bars, 10 µm (overview), 3 µm (3D-reconstruction). **(B)** Immunostaining for p62 in murine primary hippocampal neurons expressing EGFP-synphilin-1 and α-synuclein-BFP. Yellow arrows indicate p62 bodies in contact with LBLs. Scale bar, 10 µm.

**Supplementary Figure 12.**
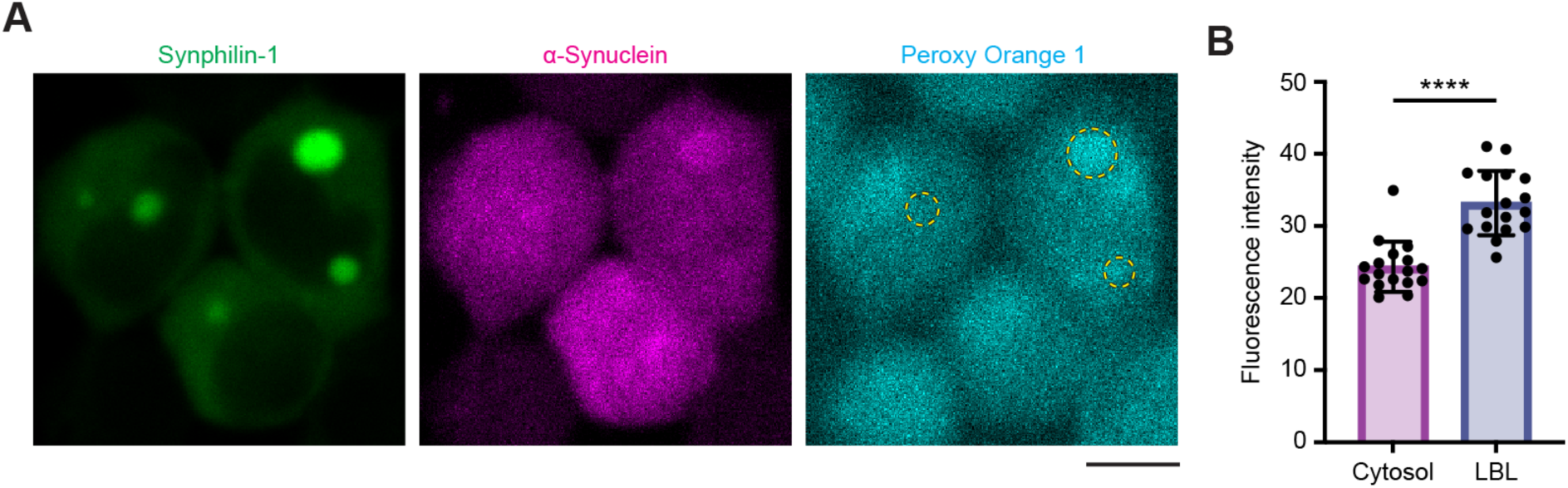
Lewy body-like condensates create a distinct redox microenvironment within cells. **(A)** Representative fluorescent images of LBLs-expressing HEK293 cells, stained with Peroxy Orange 1. Yellow dashed circles represent the location of LBLs. Scale bar, 10 µm. **(B)** LBLs maintain a distinct red/ox environment as indicated by comparison of Peroxy Orange 1 fluorescent intensity inside the condensate and in the surrounding cytosol. Data from 16 condensates, each from a different cell, from three independent replicates.

